# Accurate and Interpretable Prediction of Antidepressant Treatment Response from Receptor-informed Neuroimaging

**DOI:** 10.1101/2025.07.02.662710

**Authors:** Hanna M. Tolle, Andrea I Luppi, Timothy Lawn, Leor Roseman, David Nutt, Robin L. Carhart-Harris, Pedro A. M. Mediano

## Abstract

Conventional antidepressants show moderate efficacy in treating major depressive disorder. Psychedelic-assisted therapy holds promise, yet individual responses vary, underscoring the need for predictive tools to guide treatment selection. Here, we present `graphTRIP` (graph-based Treatment Response Interpretability and Prediction) – a geometric deep learning architecture that enables three advances: 1) accurate prediction of post-treatment depression severity using only pretreatment clinical and neuroimaging data; 2) identification of interpretable biomarkers; and 3) causal analysis of treatment effects and underlying mechanisms. Trained on data from a clinical trial comparing psilocybin and escitalopram (NCT03429075), `graphTRIP` achieves strong predictive accuracy (*r* = 0.72, *p* < 0.01) and generalises across brain atlases. Furthermore, in an independent TRD cohort, reconstruction accuracy was maintained and significant zero-shot predictions provided encouraging evidence of transferable outcome-predictive structure. The model identifies stronger functional connectivity within sensory networks as a predictor of poorer response across both treatments. In contrast, causal analysis implicates frontoparietal and default mode networks as moderators of differential response, with stronger 5-HT1A- and 5-HT2A-related signalling in the frontoparietal network predicting escitalopram response but psilocybin resistance. Overall, this work advances precision medicine and biomarker discovery in depression.

## INTRODUCTION

Major depressive disorder (MDD) is a debilitating psychiatric disorder that places a serious burden on individuals, their families, and society. Already a leading cause of disability worldwide [1], its incidence has continued to rise in recent years [2, 3], highlighting the urgent need for a more effective treatment strategy.

Selective serotonin reuptake inhibitors (SSRIs) are the most widely prescribed antidepressant drugs [4]. These compounds act by inhibiting the reuptake of serotonin (5-HT) from the synaptic cleft through blockade of the serotonin transporter (5-HTT, a.k.a. SERT) [4]. While SSRIs are among the most effective treatments currently available, less than 40% of patients achieve remission after the first treatment course, and approximately one in three fails to remit even after multiple successive treatment trials with different SSRIs [5, 6].

The psychedelic compound psilocybin has recently emerged as a promising alternative to conventional antidepressants [7], with some evidence indicating efficacy even in SSRI-resistant patients [8, 9]. Like most psychedelic drugs, psilocybin primarily acts as an agonist at the serotonin receptors 5-HT2A and 5-HT1A [10]. Notably, in contrast to SSRIs, which require continuous daily use, psilocybin treatment typically involves only one or two therapist-guided drug sessions in addition to psychological therapy [11].

However, like all treatments, psychedelic treatments come with associated risks. Individual responses to psychedelics vary widely, with one study reporting increased suicidal ideation in certain individuals [9]. This variability likely reflects both the underlying heterogeneity of MDD [12] and the marked individual differences in response to psychedelics [13]. Thus, to safely deploy psychedelic treatments and improve MDD prognosis worldwide, we need a means to predict how an individual patient will respond to a given treatment, allowing for a more targeted approach and reducing the prolonged trial- and-error process in antidepressant prescribing.

One promising approach is to predict treatment outcomes from pre-treatment neuroimaging data. Because brain function is assumed to emerge from complex inter-actions between specialised regions, it is natural to model the brain as a network, or brain graph, where nodes represent brain regions (defined by a brain atlas) and edges capture anatomical or functional connectivity [14]. Functional connectivity (FC) refers to statistical dependencies – typically correlations – between brain regional activity, and its disruption has been consistently linked to psychiatric disorders, including MDD [15–17], making it a particularly relevant feature for antidepressant response prediction. Furthermore, recent approaches such as REACT (Receptor-Enriched Analysis of functional Connectivity by Targets) enable the integration of fMRI data with normative maps of molecular targets, derived from PET imaging [18–21], providing features of neuromodulatory activity that have proven valuable for studying neurological and psychiatric disorders [22, 23], as well as the acute effects of psychedelics [24]. Together, these developments position molecularly informed brain graphs as a clinically relevant basis for predicting antidepressant treatment outcomes.

While previous studies using machine learning (ML) have achieved notable successes in predicting antidepressant treatment outcome from brain graph features [25– 28], key challenges remain for clinical translation. For instance, prior approaches have largely focused on binary classification of treatment response or remission [25, 27– 29], yet binarising continuous depression scores sacrifices valuable information and relies on statistically arbitrary thresholds. Furthermore, conventional ML architectures require fixed input sizes, which constrains the analysis to a specific brain atlas. However, the choice of atlas can substantially alter findings, and no universally accepted standard exists [30, 31].

Another key barrier to clinical translation is the lack of interpretability – most deep learning models operate as “black boxes,” offering little insight into the basis of their predictions. In a clinical context, this is undesirable. To ensure safe and reliable decisions, it is crucial to understand why a specific treatment outcome was predicted.

Finally, standard predictive approaches cannot reliably determine whether one treatment would have worked better than another for a given patient. To enable data-driven treatment selection, we must therefore move beyond mere prediction towards *causal* inference.

Here, we present a geometric deep learning (GDL) approach for predicting antidepressant treatment response from pre-treatment clinical and fMRI data in patients treated with escitalopram or psilocybin. GDL is uniquely suited to learning from brain graphs, enabling more powerful and biologically-informed predictions than conventional ML. Our novel architecture, dubbed `graphTRIP`, addresses key roadblocks to clinical translation in a fundamentally new way: 1) it directly predicts post-treatment depression scores (QIDS) with high accuracy; 2) its VGAE reconstructions generalise to an independent TRD dataset, with encouraging evidence that outcome-predictive structure also transfers; 3) it flexibly adapts to different brain atlases, maintaining significant predictions without retraining; 4) it enables rich interpretability analyses – particularly through our novel method GRAIL (Gradient Alignment for Interpreting Latent-variable models), which quantifies learned associations between treatment outcome and any brain-graph biomarkers of interest. Lastly, we extend our model within a causal inference framework [32], enabling us to estimate the expected difference in outcome under psilocybin versus escitalopram for each patient, and discern treatment-specific moderators.

Leveraging state-of-the-art GDL and causal inference, our approach advances the field of antidepressant response prediction, offering a robust tool for both biomarker discovery and clinical decision-making.

## RESULTS

The primary dataset [33] in our study included 42 MDD patients who participated in a double-blind randomised controlled trial (DB-RCT) with two treatment arms: psilocybin (*N* = 22) and escitalopram (*N* = 20) (Fig. 1a). For an out-of-distribution evaluation, we additionally used an independent dataset [8] consisting of 16 patients with treatment-resistance depression (TRD), who received psilocybin in an open-label trial (Fig. 1b).

**FIG 1.**
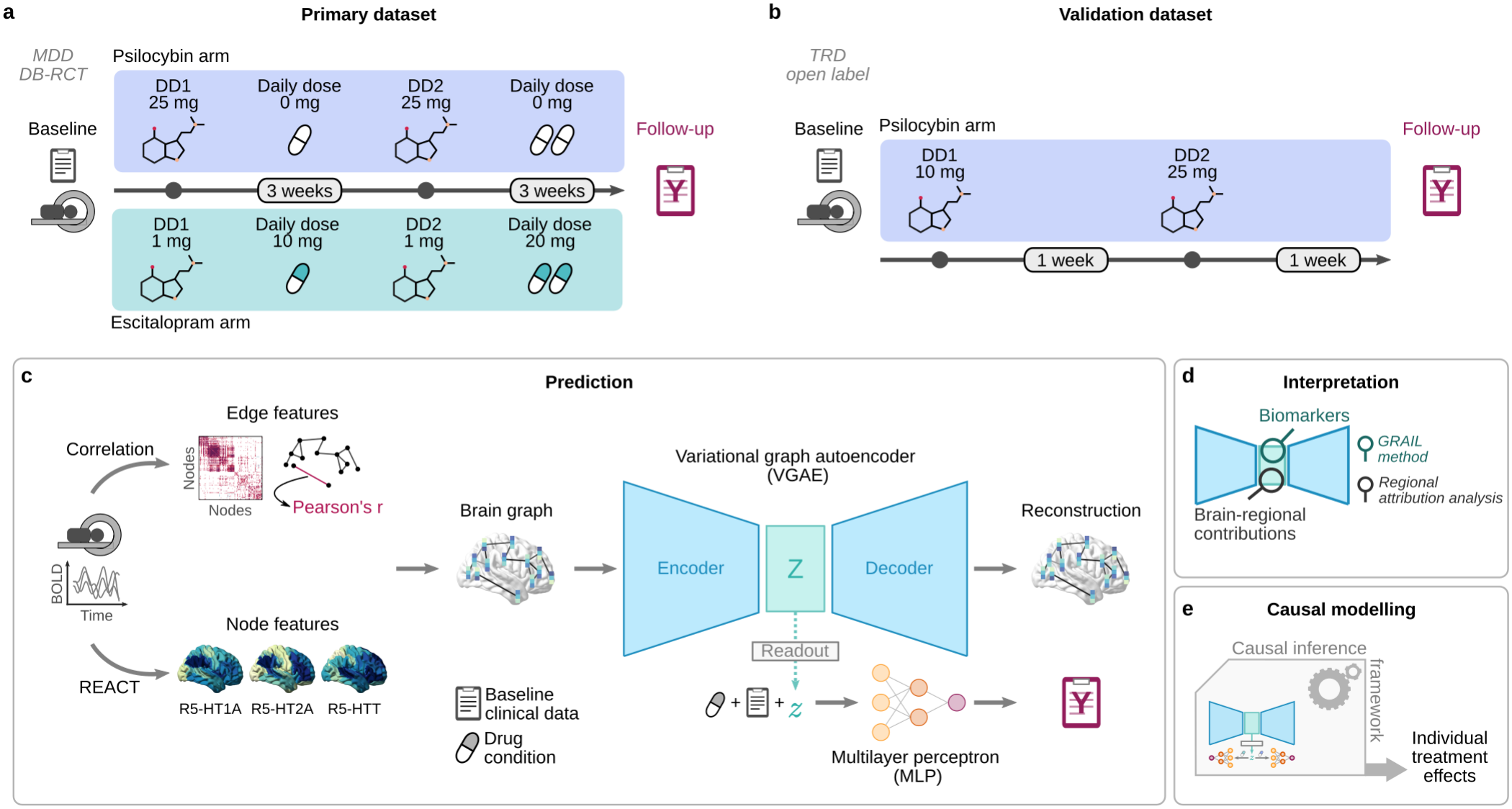
Overview of study design and methodological framework. **a**, Primary dataset: 42 MDD patients from a DB-RCT, comparing escitalopram (*N* = 20) and psilocybin (*N* = 22) treatment. Escitalopram and psilocybin groups, respectively, received inactive and high doses of psilocybin on each dosing day (DD1, DD2). Escitalopram (escitalopram arm) or placebo (psilocybin arm) was administered daily for three weeks after each session. Depression severity was assessed using QIDS three weeks after DD2. **b**, Independent dataset: 16 TRD patients received low (DD1) and high (DD2) doses of psilocybin in an open-label trial. QIDS was measured one week after DD2. **c**, Brain graphs were constructed from baseline fMRI, with node features encoding serotonin system REACT maps and edges reflecting functional connectivity. A variational graph autoencoder (VGAE) learned latent graph embeddings, which were passed to a multilayer perceptron (MLP) for predicting post-treatment QIDS. **d**, Two complementary interpretability analyses reveal the predictive contributions of brain regions and biomarkers. **e** We extend our model within a causal inference framework to estimate individual treatment effects and identify treatment-specific moderators.

Our model, called `graphTRIP` (graph-based Treatment Response Interpretability and Prediction), combines a variational graph-transformer autoencoder (VGAE) with a multilayer perceptron (MLP) (Fig. 1c). The VGAE learns a latent representation of brain graphs. The MLP predicts post-treatment QIDS. Brain graphs were constructed from baseline fMRI data, with edges defined by fixed-density thresholding of FC, retaining only the global 20% strongest edges. Node features encoded REACT values for 5-HT1A, 5-HT2A, and 5-HTT, denoted as R5-HT1A, R5-HT2A, and R5-HTT to distinguish them from the corresponding normative receptor densities. REACT provides subject-specific estimates of how strongly each brain region’s activity is functionally coupled with a molecular target of interest. The VGAE encoder maps each brain region to a latent space, and the decoder reconstructs the original graph from the resulting matrix of latent node vectors, *Z*. A readout layer then aggregates *Z* into a graph-level representation vector ***z***. This vector, combined with the drug condition (psilocybin or escitalopram) and baseline clinical data (QIDS and BDI depression scores, and a binary indicator of prior SSRI use), serves as input to the MLP.

We applied two analyses to probe model predictions: 1) Regional attribution analysis estimates brain regional contributions to the prediction; 2) GRAIL quantifies learned associations between brain-graph features and treatment outcome (Fig. 1d).

Moving beyond predictive modelling, we further extended `graphTRIP` within a causal inference framework to estimate individual treatment effects and identify treatment-specific moderators (Fig. 1e).

### Accurate, robust predictions of treatment response

`graphTRIP` achieved a strong correlation between true and predicted post-treatment QIDS scores (*r* = 0.7220; Fig. 2a) and showed good reconstruction performance (Fig. 2c-d). At the single-subject level, the median absolute prediction error was 1.96 QIDS points (IQR: 0.87-4.34; Supp. Fig. 7a). Significance was assessed against an empirical null obtained by permuting the outcome across patients and retraining the full model (Methods): none of the 100 null ensembles reached the observed value on any metric (*r, R*^2^, MAE, RMSE; *p <* 0.01, the resolution limit of 100 permutations; Fig. 2a, inset). Unless stated otherwise, all reported predictions correspond to mean test-fold predictions from 7-fold cross-validation, averaged across 10 repeated training runs with different random seeds. Full prediction metrics (*r, R*^2^, MAE, RMSE) for all models are reported in Supp. Table II. Reducing the neuroimaging data using PCA or t-SNE, or replacing the MLP with a linear output layer, failed to produce meaningful predictions, highlighting the advantage of `graphTRIP`’s architecture (Supp. Fig. 2b).

**FIG 2.**
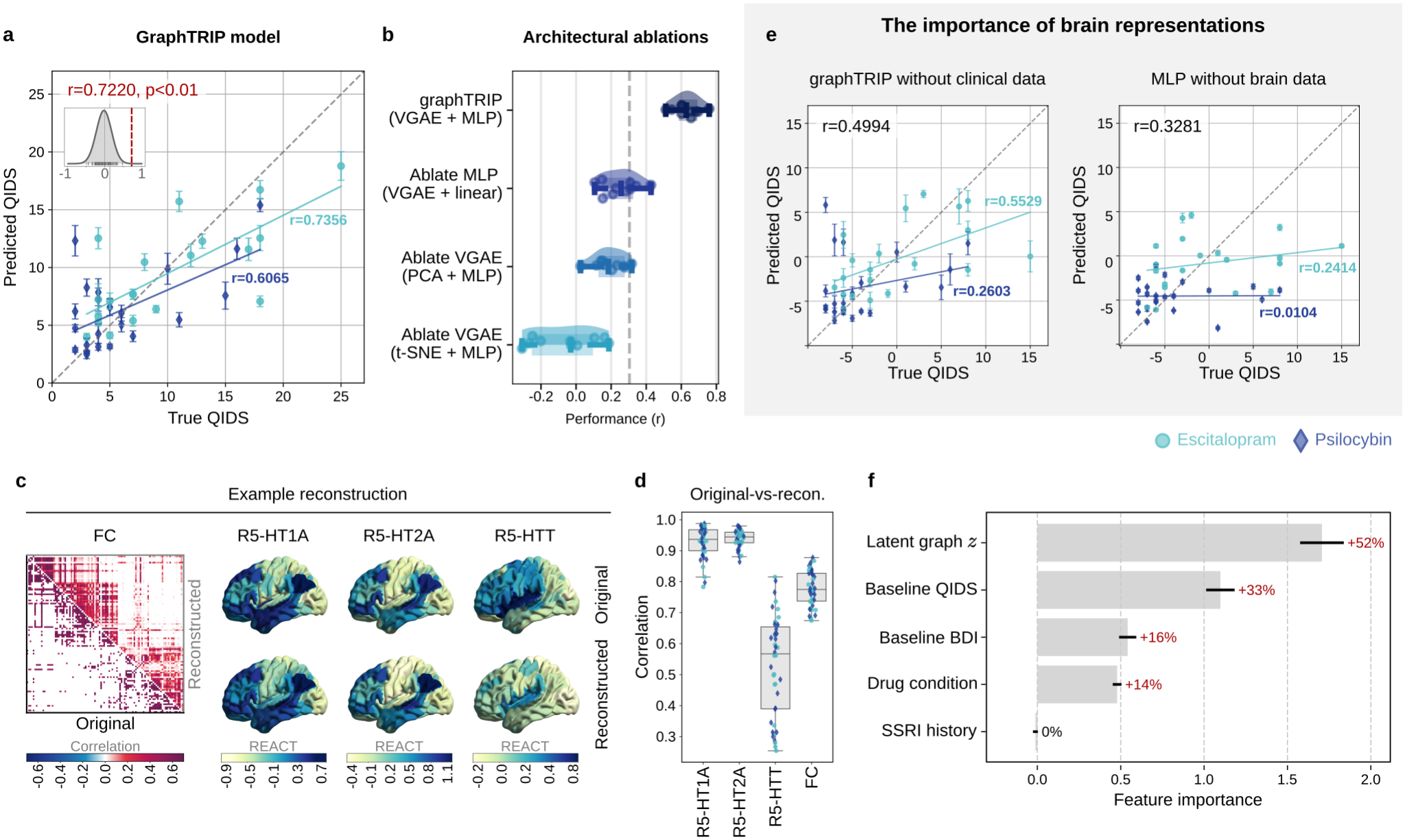
Reconstruction and prediction performance of the graphTRIP model. **a**, `graphTRIP` predicts post-treatment QIDS (*r* = 0.7220, *p <* 0.01). Inset: distribution of ensemble correlations under 100 label-permutation nulls, with the observed *r* marked. **b**, Across random seeds, `graphTRIP` outperforms architectural ablations that replace the VGAE with PCA or t-SNE, or the MLP with a linear output layer. **c**, Example VGAE reconstructions of FC and REACT node features. **d**, Correlations between original and reconstructed FC and node features across all patients. **e**, A `graphTRIP` model trained on brain graphs and drug condition without clinical data still predicts outcome (left); an MLP trained only on clinical data and drug condition performs substantially worse (right). **f**, Permutation importance analysis, showing the increase in mean absolute error (MAE) caused by shuffling individual features across patients. Intuitively, shuffling important features results in greater prediction error. Bars indicate the mean increase in MAE across 50 random permutations; error bars denote standard error. Percentage annotations report this increase relative to each model’s per-seed out-of-fold baseline MAE.

To assess the relative contribution of brain and clinical features, we trained ablated models on each input type alone. A brain-only `graphTRIP` model (brain graphs and drug condition, without clinical data) still predicted post-treatment QIDS (*r* = 0.4994; Fig. 2e), whereas a clinical-only MLP with the same prediction-head architecture performed worse (*r* = 0.3281; Fig. 2e). Despite its moderate pooled correlation, the clinical-only model showed weak within-arm associations in both groups (psilocy-bin: *r* = 0.0104; escitalopram: *r* = 0.2414), suggesting that it primarily predicted each treatment arm’s mean post-treatment QIDS, which was higher under escitalopram [33]. The brain-only model improved within-arm prediction substantially (psilocybin: *r* = 0.2603; escitalopram: *r* = 0.5529), and the full `graphTRIP` model was strongest in both arms (psilocybin: *r* = 0.6065; escitalopram: *r* = 0.7356).

These results suggest that brain features contribute most to outcome prediction. Several follow-up analyses corroborate this: First, permutation importance analysis measured the impact of shuffling each input feature on prediction error (mean absolute error) across 50 repetitions, with larger increases indicating greater importance. The latent brain-graph feature vector ***z*** (32 dimensions, shuffled as a group) was the most important predictor, followed by baseline QIDS and BDI scores (Fig. 2f). The contribution of baseline QIDS is consistent with its significant correlation with post-treatment QIDS (*r* = 0.3981, *p* = 0.0090), although this association differed between treatment arms (escitalopram: *r* = 0.5297, *p* = 0.0163; psilocybin: *r* = − 0.0027, *p* = 0.9905). Second, partial correlation analysis showed that the association between true and predicted `graphTRIP` scores remained strong after controlling for drug condition, baseline QIDS, and their interaction (*r* = 0.6284). Finally, linear probes and MLP prediction heads trained only on frozen VGAE representations and the drug condition still yielded strong predictions (linear: *r* = 0.5435; MLP: *r* = 0.6072). Together, these converging results imply that the neuroimaging data contains outcome-relevant information, and that `graphTRIP` was able to extract this information.

We additionally trained a `graphTRIP` model to predict post-treatment BDI scores (*r* = 0.5254; Supp. Fig. 8), confirming the robustness of our approach.

### Generalisation across brain atlases

We evaluated `graphTRIP`’s ability to generalise across brain parcellations by testing it – without retraining– on atlases different from the one used for training (Schaefer 100 [34]). This is possible because the VGAE shares parameters across nodes and pools them into a fixed-size graph embedding, so neither the encoder nor the MLP depends on the number of parcels (see Methods). On Schaefer 200, which has twice as many parcels, `graphTRIP` maintained highly accurate reconstructions and QIDS prediction performance (*r* = 0.7156, *p <* 0.01; Fig. 3), comparable to performance on Schae-fer 100 (*r* = 0.7220). We further replicated this result on the AAL atlas with 116 regions [35], which differs fundamentally from Schaefer and includes subcortical regions, where `graphTRIP` also generalised well (*r* = 0.6049, *p <* 0.01; Supp. Fig. 9).

**FIG 3.**
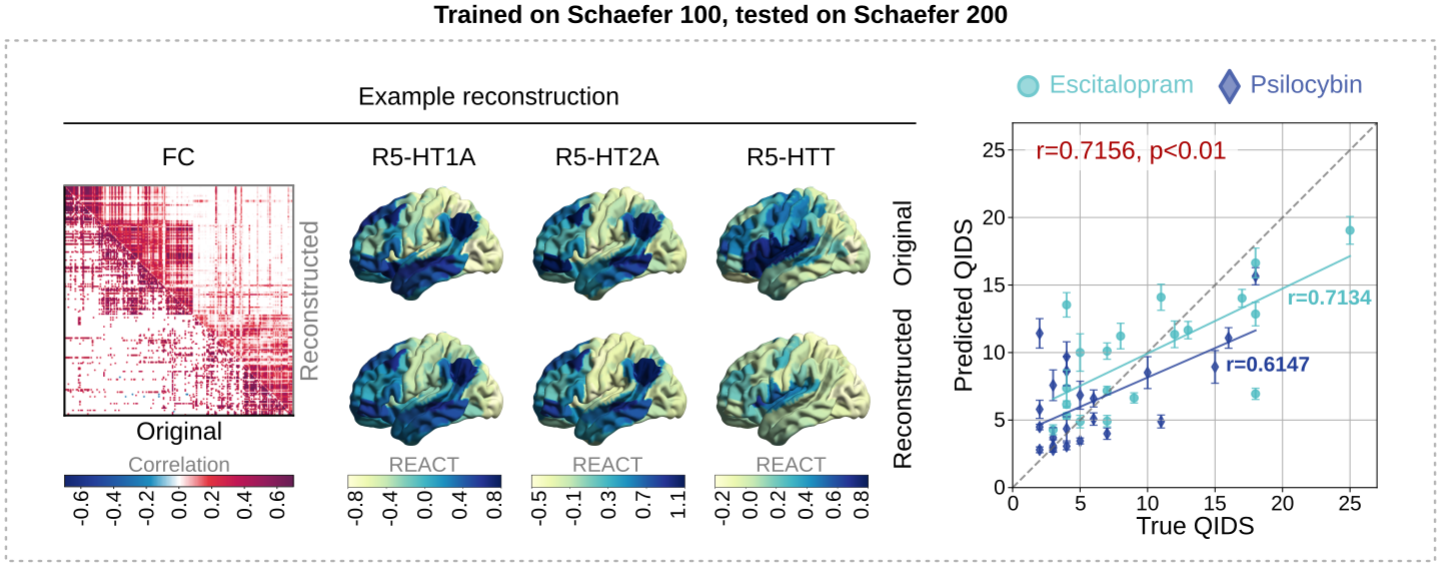
graphTRIP generalises across brain atlases without retraining. The `graphTRIP` model, trained on the Schaefer 100 brain atlas, generalises to Schaefer 200 while maintaining strong reconstruction and prediction performance. The panel shows FC and node feature reconstructions for an example subject, and the MLP’s prediction performance.

### Evaluation on an independent dataset

We evaluated `graphTRIP` on an independent dataset that differed from the primary dataset in several key aspects (Fig. 4a), including the presence of only one treatment arm (psilocybin), a different psilocybin treatment protocol, patients with treatment-resistant depression (TRD), and a different follow-up time point. Patients in the independent dataset also had substantially higher baseline QIDS and BDI scores (Fig. 4b).

**FIG 4.**
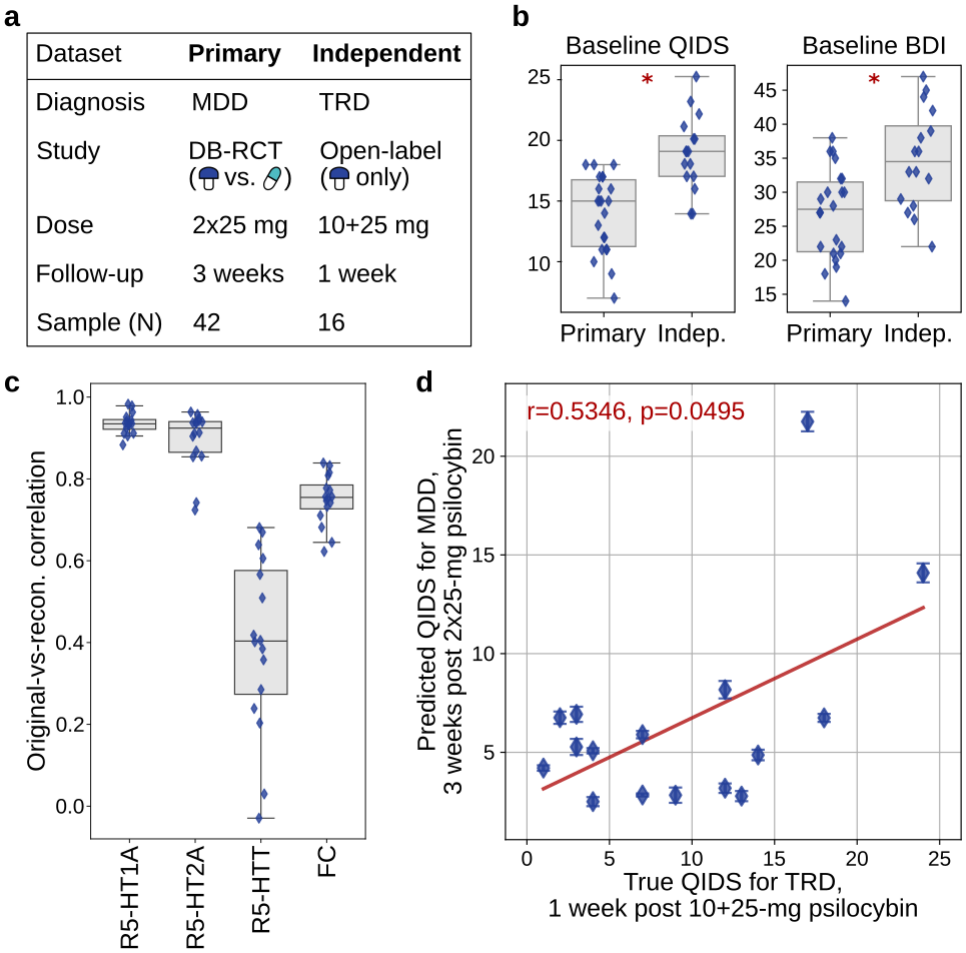
Zero-shot evaluation on an independent TRD dataset. **a**, Key differences between the primary (MDD, DB-RCT) and independent (TRD, open-label) datasets. **b**, Baseline QIDS and BDI in the primary versus independent datasets, indicating that the model is asked to extrapolate to significantly more severely depressed patients. **c**, Original versus reconstructed brain-graph feature correlations on the independent dataset (VGAE applied without retraining). **d**, Zero-shot QIDS prediction on the independent TRD dataset (baseline clinical inputs rescaled onto the training scale; model not retrained).

Despite these differences, VGAE reconstruction accuracy was maintained without retraining: the reduction relative to the primary dataset was modest and statistically non-significant (Fig.4c; unpaired Mann-Whitney U tests on per-subject originalfig-vs-reconstructed correlations vs. Fig.2d: all FDR-corrected *p >* 0.05). Because 9/16 patients from the independent dataset had base-line QIDS higher than any psilocybin-arm patient in the training dataset (Fig.4b), baseline QIDS and BDI were renormalized to each fold model’s training input scale before evaluating QIDS predictions (Methods); brain-graph inputs and model weights were unchanged. Under this protocol, `graphTRIP`’s zero-shot predictions were significantly correlated with observed QIDS in the independent cohort (*r* = 0.5346, permutation *p* = 0.0495; Fig.4d).

In the independent dataset, baseline QIDS was significantly associated with one-week post-treatment QIDS (*r* = 0.5729, *p* = 0.0204), and the clinical-only MLP from Fig.2e, evaluated under the same protocol, achieved a similar zero-shot correlation as `graphTRIP` (*r* = 0.5271). However, in the primary-dataset psilocybin arm, baseline and 3-weeks post-treatment QIDS were uncorrelated (*r* = −0.0027, *p* = 0.9905), and in the same TRD cohort of the independent dataset, the baseline-outcome association was absent at three-month follow-up (*r* = 0.2307, *p* = 0.3901). This suggests that a predictor based solely on baseline severity might not generalise. An important question is therefore whether `graphTRIP`’s latent brain features contribute predictive signal beyond baseline severity. Permutation importance analysis provides supporting evidence: shuffling brain-graph features across patients reduced `graphTRIP`’s zero-shot correlation by 0.1872 on average; shuffling baseline QIDS reduced it by 0.3763. In contrast, shuffling baseline QIDS for the clinical-only MLP collapsed predictive performance entirely (mean drop in *r* = 0.6128).

These results provide encouraging evidence that the learned brain representation and outcome-predictive structure transfer across cohorts.

### Estimating individual treatment effects

Identifying which treatment is most effective for a given patient requires estimating individual treatment effects (ITEs) in a causal analysis. This relies on several assumptions, including strong ignorability (no unmeasured confounding given observed covariates). RCTs are designed to approximate these conditions, and we therefore assume that they largely hold for the present analysis. We note, however, that small-sample effects and post-randomisation factors such as dropouts can introduce subtle imbalances. Indeed, we observed an almost-significant difference in baseline QIDS between treatment groups, with lower baseline severity in the psilocybin arm (two-sample *t*-test: *t* = 1.9928, *p* = 0.0531, Cohen’s *d* = 0.6104).

Under these assumptions, one could in principle estimate ITEs using `graphTRIP` by generating predictions for each patient under both treatment conditions. However, while `graphTRIP` provides a robust predictor of post-treatment QIDS, its predictions under psilocybin and escitalopram for the same subject were almost perfectly correlated and differed primarily by a near-constant off-set (Supp. Fig. 10). This suggests that the model captured an average treatment effect but did not learn strong brain-drug interaction effects. This behaviour is consistent with known limitations of single-model approaches like `graphTRIP`, which can underfit treatment-covariate interactions and thus attenuate heterogeneous treatment effects when baseline features are substantially more predictive of outcomes than the treatment itself [36].

To explicitly model potential drug-specific response patterns, we developed `Medusa-graphTRIP` – an extension of `graphTRIP` inspired by the Counterfactual Regression framework [32]. `Medusa-graphTRIP` uses a shared VGAE to learn a common latent brain-graph representation across both treatment arms, followed by two separate MLP heads that predict post-treatment QIDS, one for each treatment condition (Fig. 5a).

**FIG 5.**
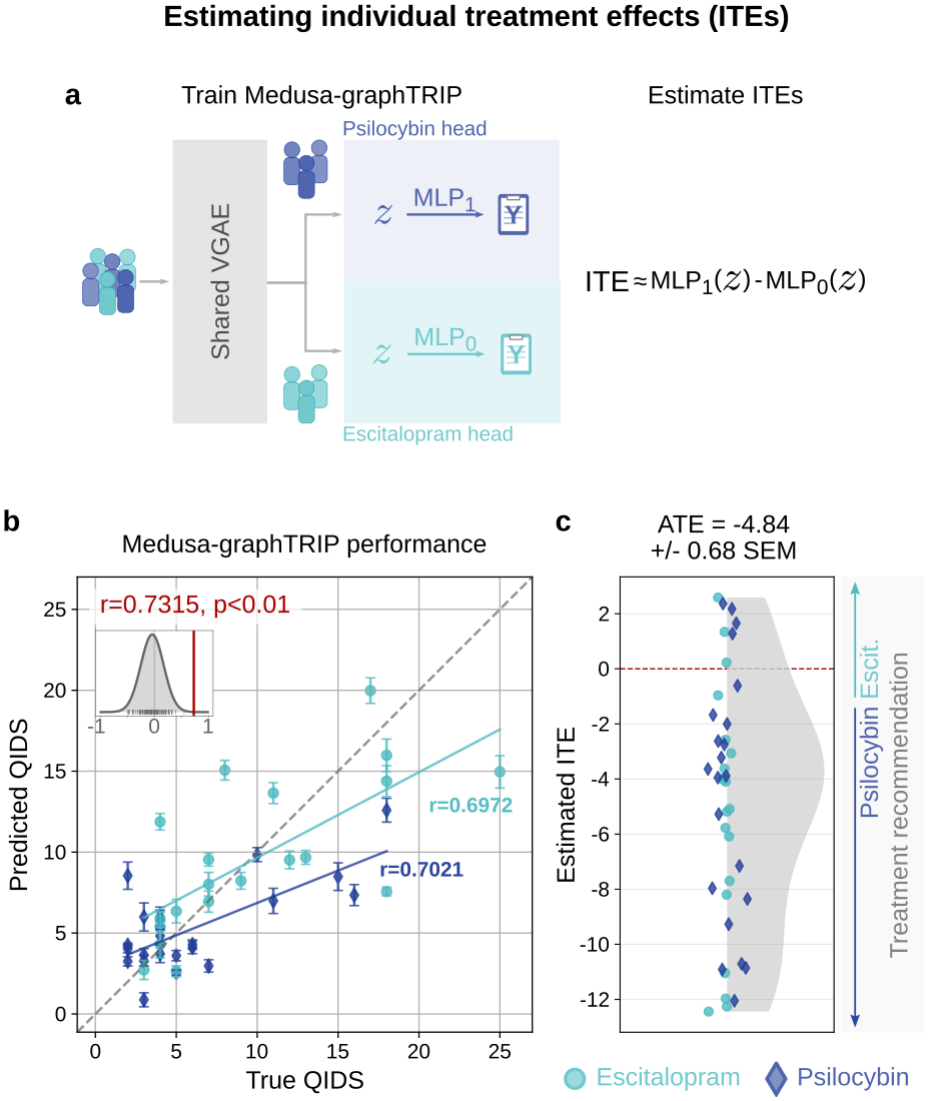
Causal inference with Medusa-graphTRIP enables estimation of individual treatment effects (ITEs). **a**, Schematic of the `Medusa-graphTRIP` architecture. A shared VGAE learns a common latent representation of pre-treatment brain graphs, which is passed to two separate MLP outcome heads trained on data from a single treatment condition each (psilocybin, MLP_1_; escitalopram, MLP_0_). ITEs are estimated as the difference between predictions from the two heads. **b**, `Medusa-graphTRIP` predicts post-treatment QIDS (*r* = 0.7315, *p <* 0.01). Inset: distribution of ensemble correlations under 100 label-permutation nulls, with the observed *r* marked. **c**, Estimated individual treatment effects indicate that most patients are predicted to respond better to psilocybin than escitalopram, with an average treatment effect (ATE) of −4.84 *±* 0.68 standard error.

Compared to a single-head model, this two-head structure tends to be better at capturing heterogeneous treatment effects when response functions differ across treatments [36]. However, since each head is trained on fewer samples, this approach trades reduced bias for increased variance. Thus, we use `graphTRIP` as a more stable estimator of general treatment response, whereas `Medusa-graphTRIP` is introduced to disentangle treatment-specific response patterns.

`Medusa-graphTRIP` predicted post-treatment QIDS with high accuracy (*r* = 0.7315, *p <* 0.01 vs. label-permutation null; Fig. 5b), and produced faithful brain-graph reconstructions (Supp. Fig. 11a). Its median absolute prediction error was 2.07 QIDS points (IQR: 1.28– 3.54; Supp. Fig. 7b). ITEs were estimated as the difference between predictions from the psilocybin and escitalopram heads, such that negative values indicate predicted better response to psilocybin (Fig. 5a). Consistent with the significant group difference in post-treatment QIDS in the data (two-sample *t*-test: *t* = 2.2174, *p* = 0.0323, Cohen’s *d* = 0.6805), most patients were predicted to respond better to psilocybin (Fig. 5c), with an estimated average treatment effect of ATE = −4.84 *±*0.68 (standard error). Nevertheless, seven patients were estimated to benefit more from escitalopram.

### Relative contributions of FC and REACT

We assessed the relative contributions of FC and REACT node features to prediction performance on `graphTRIP` and `Medusa-graphTRIP` by training ablated variants of each model in which one or more input domains were removed, keeping everything else unchanged (Supp. Fig. 12; Supp. Table II). All `graphTRIP` variants used drug condition as input. We compared: the full model (FC, REACT, and clinical covariates); FC with clinical covariates (REACT removed); FC with REACT (clinical removed); and FC only (REACT and clinical removed). All of these variants, including FC only, still included the conditional node features that encode each region’s MNI coordinates (see Methods).

All ablations degraded `graphTRIP`’s performance (full model: *r* = 0.7220; FC and clinical: 0.6431; FC and REACT: 0.4994; FC only: 0.3931), though the effect of removing REACT did not reach significance in a Wilcoxon test at ten seed pairs. For `Medusa-graphTRIP`, every ablation reduced performance significantly (full model: *r* = 0.7315; FC and clinical: 0.5483; FC and REACT: 0.6233; FC only: 0.5595; all *p <* 0.0040 after FDR-correction). Overall, these results suggest that while `graphTRIP`’s predictions predominantly rely on FC, `Medusa-graphTRIP` critically relies on both FC and REACT.

### Regional attributions align with the unimodal-transmodal cortical hierarchy

We assessed the contributions of individual brain regions to predicting 1) post-treatment QIDS using `graphTRIP`, and 2) ITEs using `Medusa-graphTRIP`. To this end, we performed a regional attribution analysis, which quantifies how small perturbations to each region’s latent representation influences the predicted outcome for a given patient. In contrast to the input-domain ablations, regional attributions measure the predictive contributions of latent brain-region representations, which encode both FC and REACT inputs. Specifically, the gradient of the predicted outcome *ŷ* with respect to the latent space decomposes into node-wise partial gradients 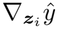, one for each brain region *i*. Regional attribution scores are defined as the percentage energy (sum of squares) of each node-wise gradient from the total energy of the full gradient vector. This gradient-based approach is conceptually similar to established attribution methods [37].

Averaging attribution profiles across patients revealed that `graphTRIP` assigns the largest contributions to sensory-motor (SMN) and visual (VIS) regions, and the smallest contributions to association cortices (Fig. 6a). In contrast, `Medusa-graphTRIP`’s ITE estimates relied most strongly on regions within the default mode (DMN) and frontoparietal (FPN) networks, with comparatively lower contributions from SMN and VIS regions (Fig. 6b). These mirror-reversed spatial patterns closely align with the unimodal-transmodal cortical gradient, which captures key anatomical, functional, and neurochemical distinctions across the cortex [20, 38–41].

**FIG 6.**
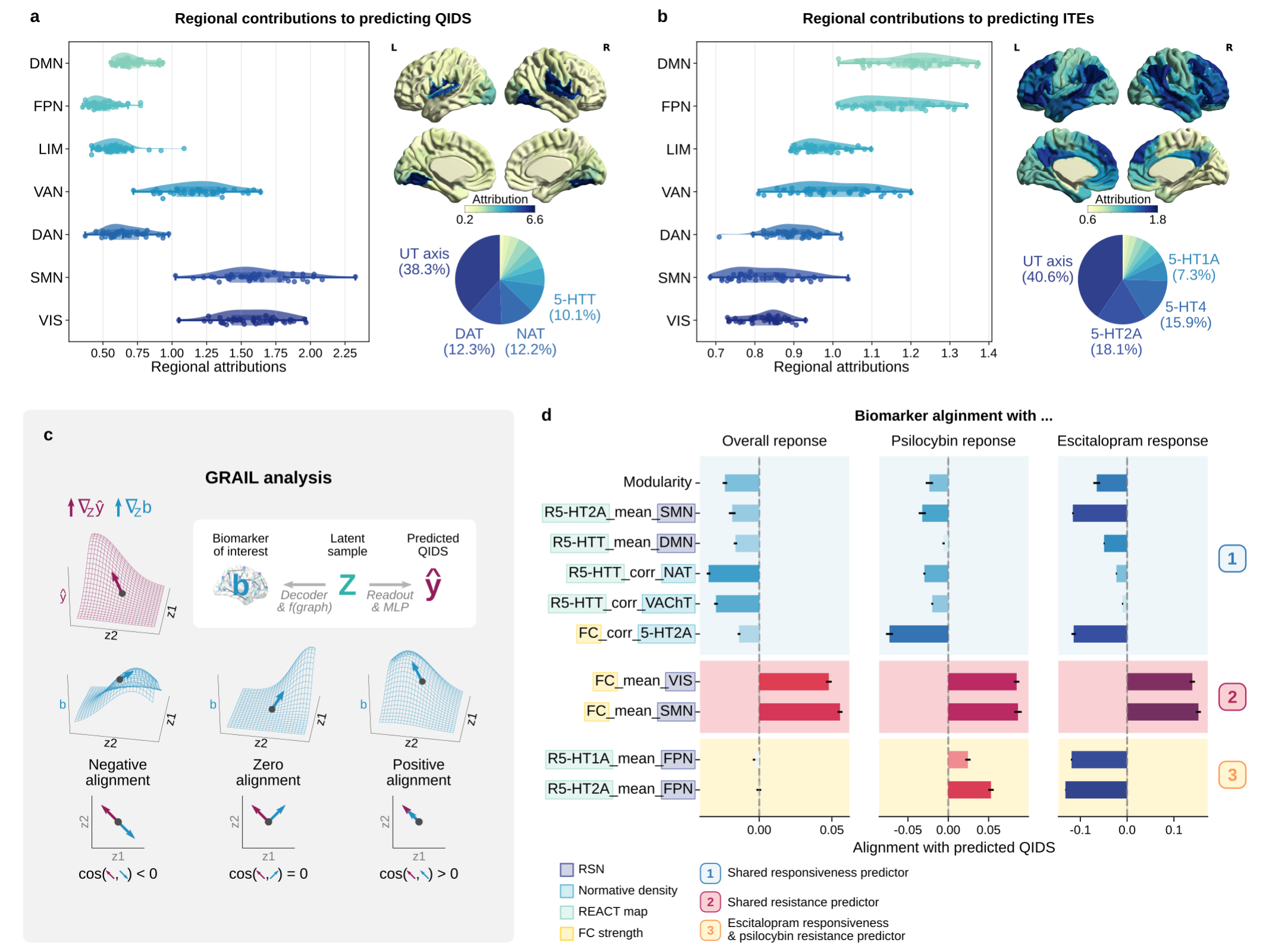
Interpretability analyses reveal unimodal regions as general outcome predictors and transmodal cortices as treatment-specific moderators. **a**, Regional attribution analysis for `graphTRIP` quantifies each brain region’s contribution to predicted post-treatment QIDS. Left: mean attributions aggregated within resting-state networks (RSNs). Top right: population-mean regional map. Bottom right: dominance analysis from a linear model explaining regional attributions from unimodal–transmodal (UT) axis scores and normative neuroreceptor density maps; pie segments indicate each regressor’s relative contribution to explained variance (*R*^2^). **b**, Regional attribution analysis applied to `Medusa-graphTRIP` individual treatment effect (ITE) predictions. **c**, Overview of GRAIL, which estimates associations between candidate biomarkers and model predictions via gradient alignment in latent space. **d**, Biomarker alignments with the predictions of three models: `graphTRIP` (left), and `Medusa-graphTRIP`’s psilocybin head (centre) and escitalopram head (right). Positive/negative values indicate association with worse/better outcomes under both drugs, psilocybin, and escitalopram, respectively. Bars show mean *±* standard error across patients. Numbers indicate biomarker categorizations based on alignment directions across models.

**FIG 7.**
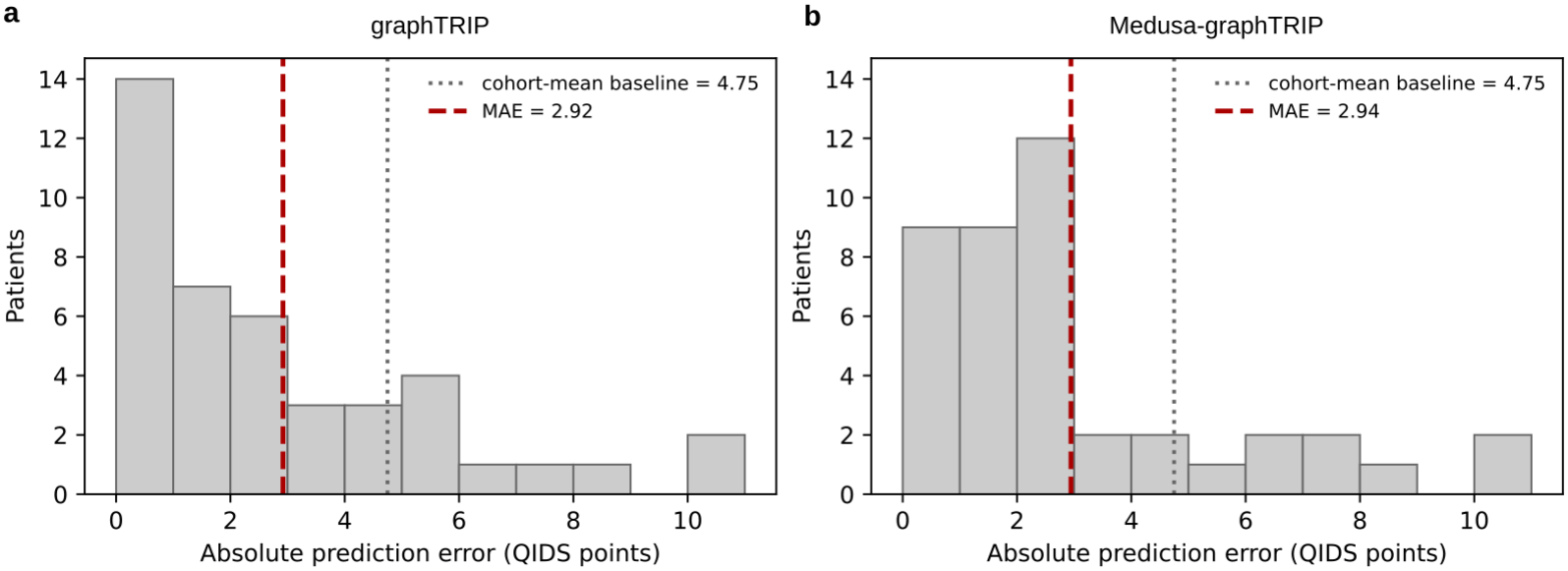
Distribution of single-subject absolute prediction errors. Absolute differences between observed post-treatment QIDS and mean out-of-fold predictions averaged across ten random seeds for **a**, `graphTRIP`, and **b**, `Medusa-graphTRIP`. Dashed red lines show the mean absolute error (MAE); dotted grey lines show the MAE that we would obtain by simply predicting the cohort-mean outcome for every patient.

**FIG 8.**
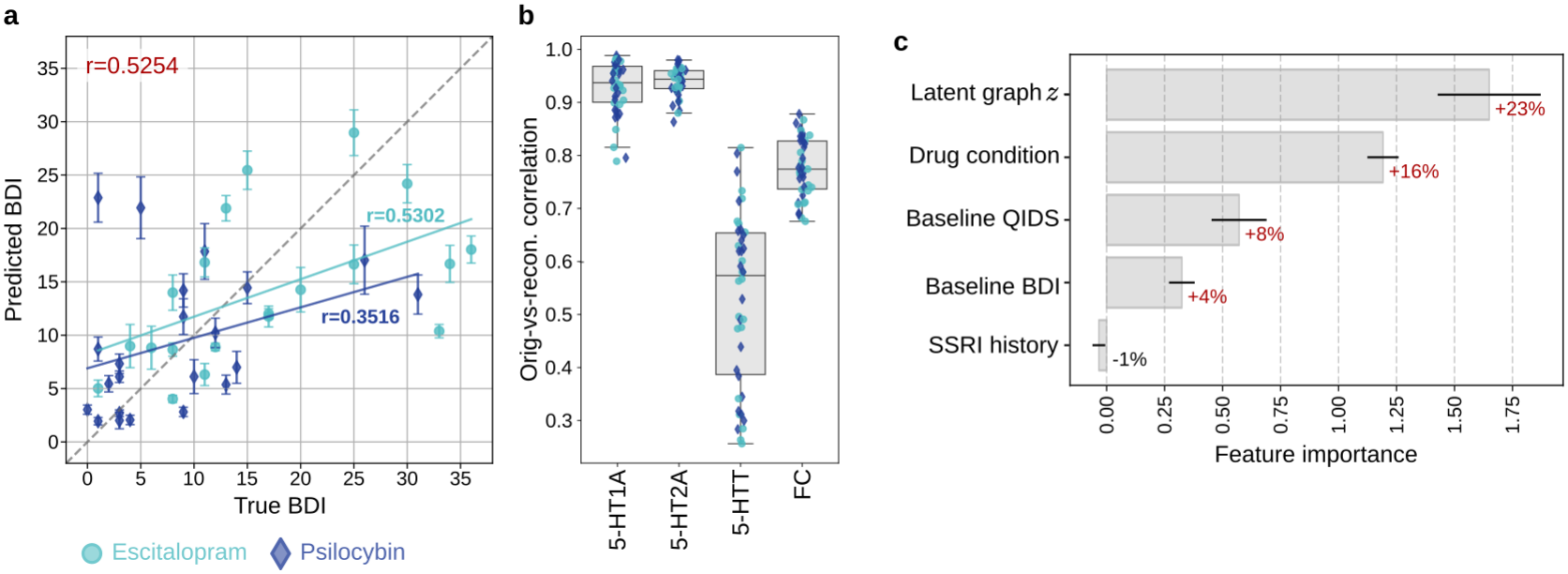
Predicting post-treatment BDI scores with graphTRIP. A separate `graphTRIP` model was trained to predict post-treatment BDI scores, using the same configuration and training parameters as the main model predicting post-treatment QIDS. This analysis demonstrates the flexibility of our pipeline for alternative clinical outcomes. **a**, True versus predicted BDI scores. **b**, Correlations between original and reconstructed edge and node features confirm accurate graph reconstruction by the VGAE. **c**, Permutation importance analysis reveals that latent brain-graph features are the most influential predictors of BDI scores. Bars indicate the mean increase in mean absolute error (MAE) across 50 random permutations when a given feature is shuffled across patients; error bars denote standard error. Percentage annotations report this increase relative to each model’s per-seed out-of-fold baseline MAE.

**FIG 9.**
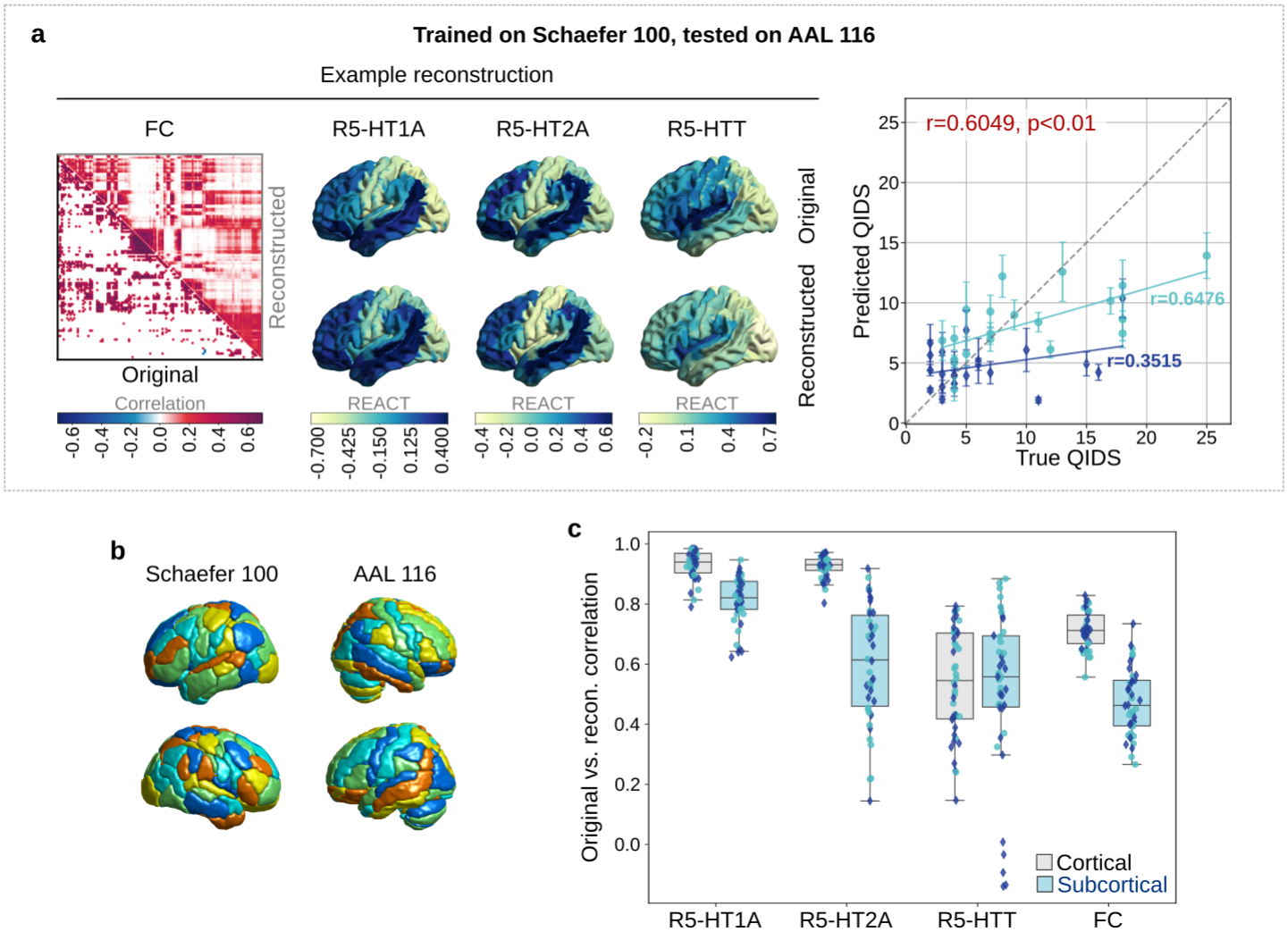
graphTRIP generalises to brain graphs defined using the AAL atlas. **a**, `graphTRIP`, trained on brain graphs constructed with the Schaefer 100 atlas, generalises to graphs using the AAL atlas, maintaining strong reconstruction and prediction performance. The panel shows example reconstructions of FC edges (left) and node features (center), and QIDS prediction performance (right). **b**, Brain renderings of the Schaefer 100 and AAL 116 atlases, comprising 100 and 116 brain regions, respectively. **c**, Correlations between original and reconstructed edge and node features for all patients. The AAL atlas includes subcortical regions not present in the Schaefer atlas. To assess model performance separately for these, reconstruction correlations are shown for cortical and subcortical regions. As expected, reconstruction is slightly weaker for subcortical regions, which the model never encountered during training, but remains clearly positive.

**FIG 10.**
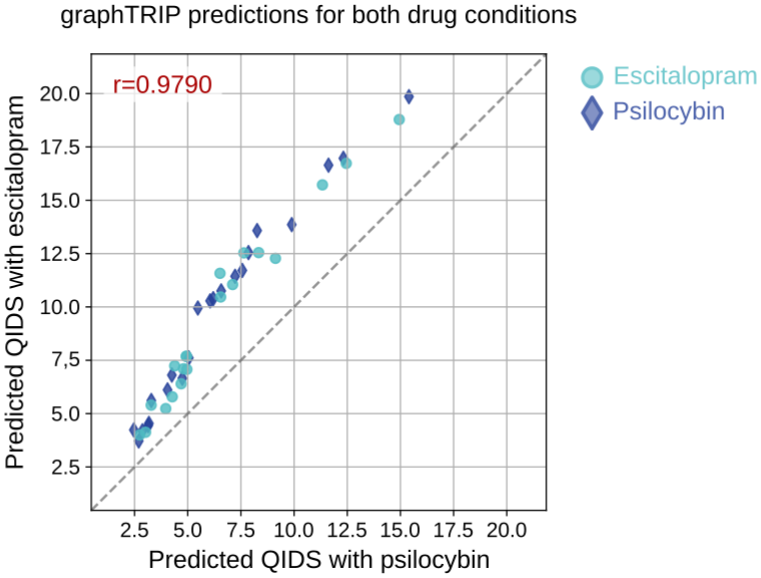
graphTRIP shows limited sensitivity to treatment condition. Scatter plot of `graphTRIP` predictions for each patient when the treatment indicator is fixed to escitalopram (y-axis) versus psilocybin (x-axis). Predictions are nearly perfectly correlated and differ primarily by a near-constant offset, indicating that `graphTRIP` captures strong treatment-invariant prognostic signal while only weakly modelling treatment-specific interactions.

**FIG 11.**
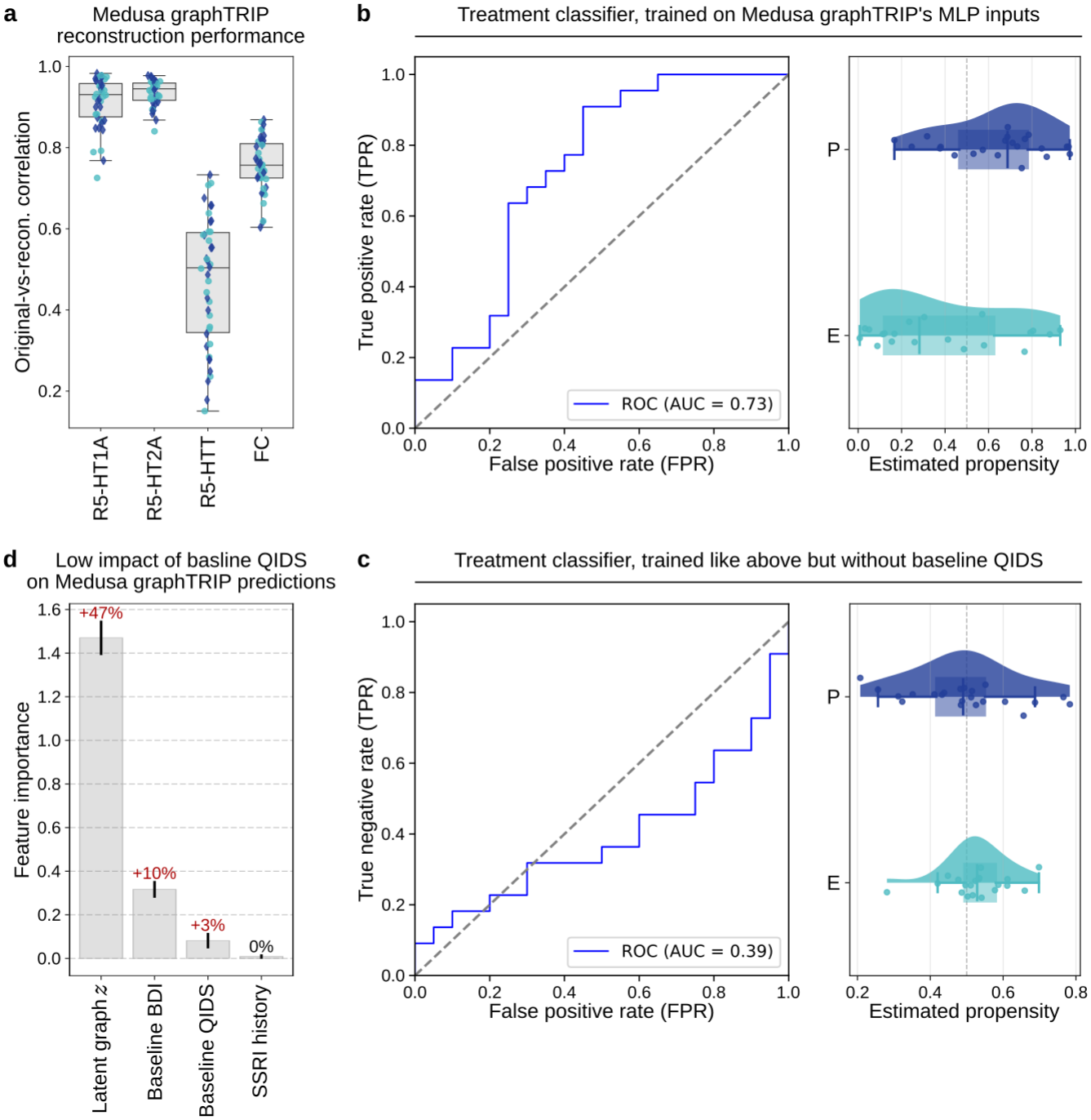
Model validation and overlap assessment for Medusa-graphTRIP. **a**, Correlations between original and recon-structed brain-graph features indicate good reconstruction performance of `Medusa-graphTRIP`’s VGAE. **b**, Treatment assignment was modestly predictable from pre-treatment covariates (AUC= 0.73). A logistic-regression model trained on the inputs to the `Medusa-graphTRIP` outcome heads (frozen VGAE latent representations and clinical covariates) achieved above-chance classification of treatment groups. The ROC curve (left) and the distribution of predicted treatment probabilities (right) indicate that, despite this predictability, estimated propensities remain bounded away from 0 and 1 for most patients, suggesting sufficient overlap between treatment groups. **c**, When baseline QIDS was excluded from the classifier inputs, treatment prediction performance collapsed (AUC= 0.39), with predicted probabilities concentrated near 0.5 for both treatment conditions, indicating that treatment predictability was driven primarily by baseline severity. **d**, Permutation importance analysis for `Medusa-graphTRIP` outcome prediction shows that latent brain-graph representations contribute most strongly to model predictions, whereas baseline QIDS exhibits only weak influence. Bars indicate the mean increase in mean absolute error (MAE) across 50 random permutations; error bars denote standard error. Percentage annotations report this increase relative to each model’s per-seed out-of-fold baseline MAE.

**FIG 12.**
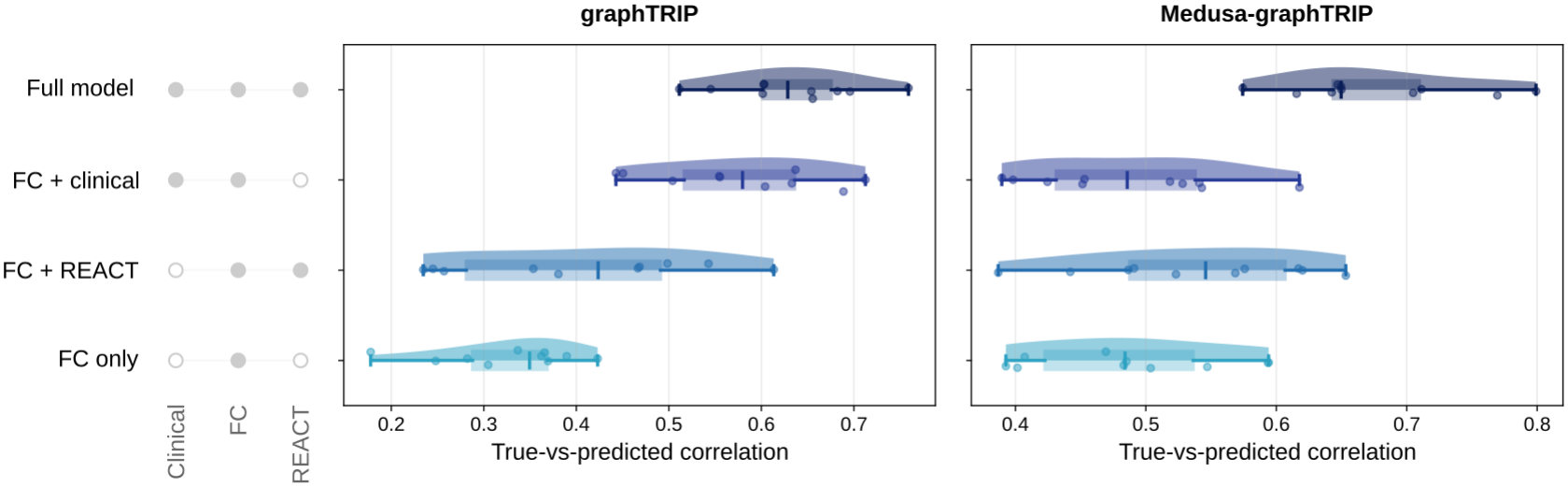
Input-domain ablations for graphTRIP and Medusa-graphTRIP. Distribution of test-fold correlations between true and predicted post-treatment QIDS across 10 repeated training runs with different random seeds, for ablation models with specific input features removed (everything else unchanged compared to the core model). Models are ordered from the full model (top) to FC only (bottom). All `graphTRIP` variants retain drug condition as an input; `Medusa-graphTRIP` uses drug condition only to route prediction through the treatment-specific heads.

**FIG 13.**
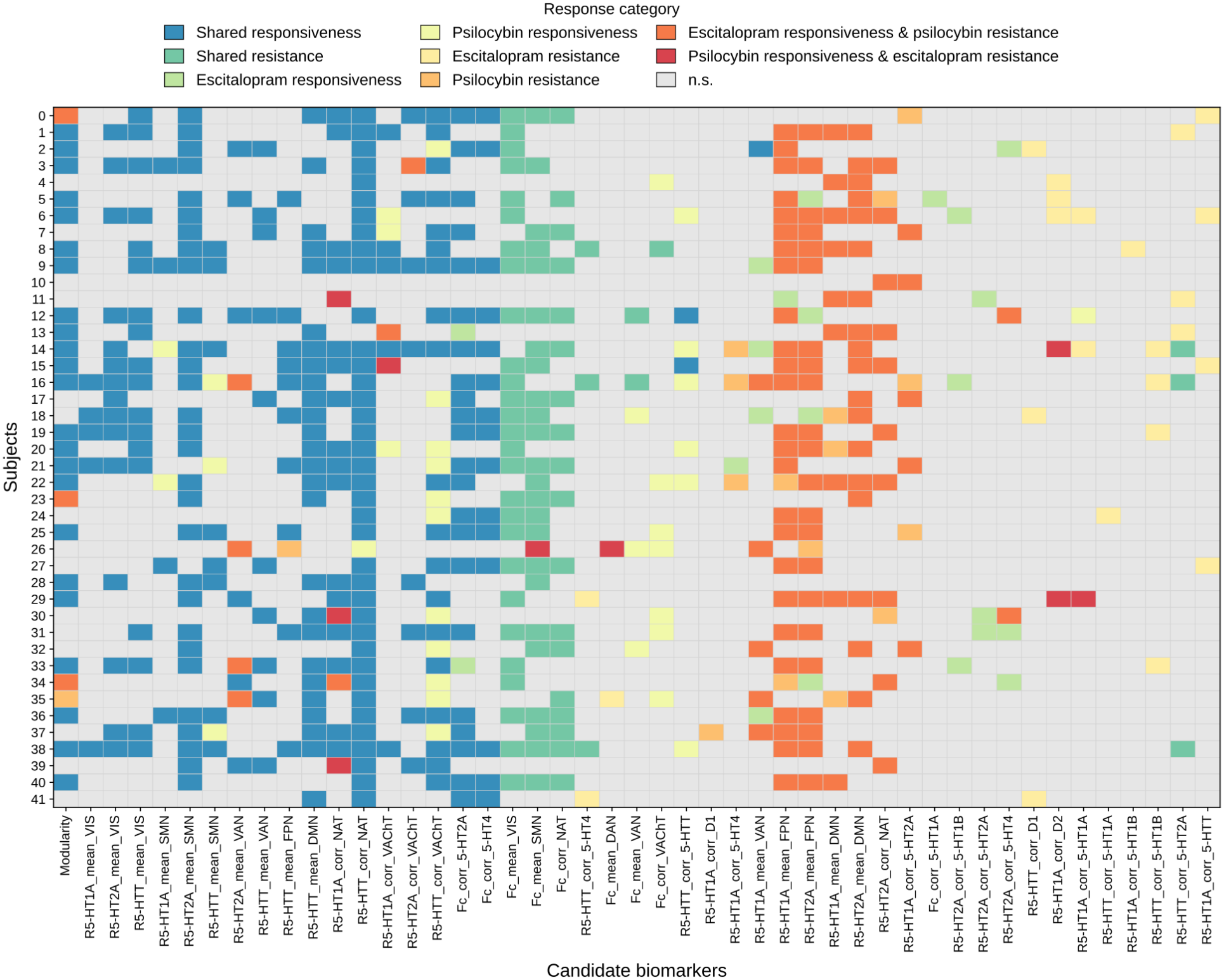
Per-patient GRAIL biomarker category assignments. Heatmap of category assignments for each subject (rows) and candidate biomarker (columns). Colours indicate the assigned biomarker category based on significant alignment directions across GRAIL modes (Table I); grey cells indicate non-significant or invalid assignments.

**FIG 14.**
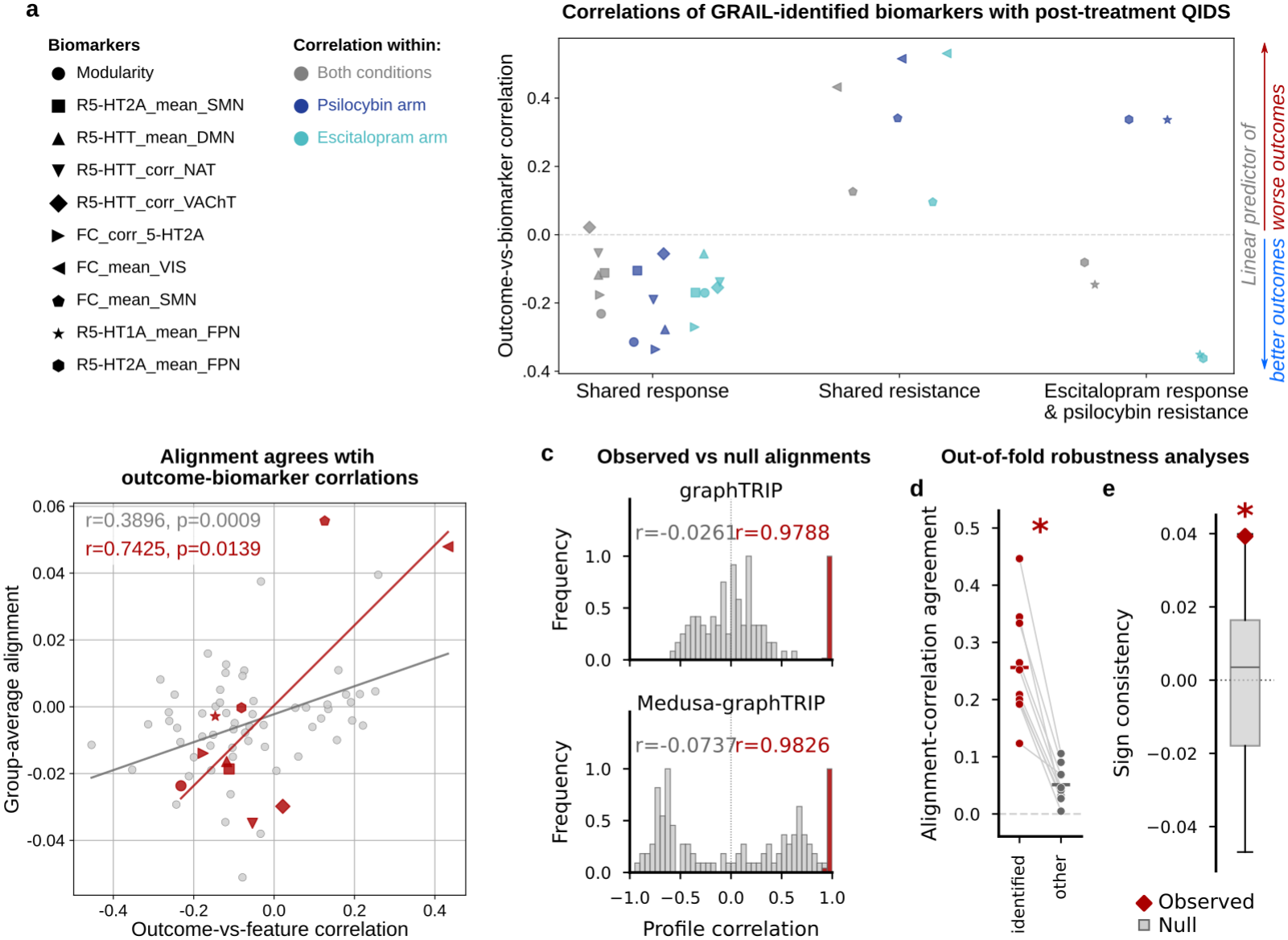
Corroboration of GRAIL biomarker associations. **a**, Univariate correlations between each GRAIL-identified biomarker and post-treatment QIDS. Grey markers show correlations computed across all patients, while blue and green markers show within-condition correlations for the psilocybin and escitalopram groups, respectively. Biomarkers are grouped by the category assigned by our analysis, illustrating that the direction of univariate associations matches the direction of GRAIL alignment: shared responsiveness predictors show negative correlations with outcome, shared resistance predictors show positive correlations, and biomarkers classified as escitalopram responsiveness but psilocybin resistance predictors show opposite correlation signs across treatment arms. **b**, Correspondence between group-mean GRAIL alignment and Pearson biomarker– outcome correlations across all 69 candidate biomarkers (*r* = 0.3896, *p* = 9.4 *×* 10^−4^) and across the ten identified biomarkers (*r* = 0.7425, *p* = 0.0139). **c**, Reproducibility of patient-by-biomarker GRAIL profiles. Profiles were highly correlated between split halves of the observed-outcome seeds (graphTRIP: mean *r* = 0.9788; `Medusa-graphTRIP`: mean *r* = 0.9826), but not between observed-outcome and label-permutation models (mean *r* = −0.0261 and −0.0737, respectively). **d**, Out-of-fold correspondence between training-set GRAIL alignments and biomarker-outcome correlations in held-out patients. Correspondence was greater for the identified biomarkers than for the remaining candidates (one-sided paired Wilcoxon test across ten seeds, *p* = 0.0010). **e**, Out-of-fold sign agreement across all 69 candidate biomarkers, without biomarker selection. Training-set GRAIL alignment agreed with held-out biomarker-outcome associations significantly better for `graphTRIP` than for label-permutation models (one-sided permutation test, *p* = 0.0198).

**FIG 15.**
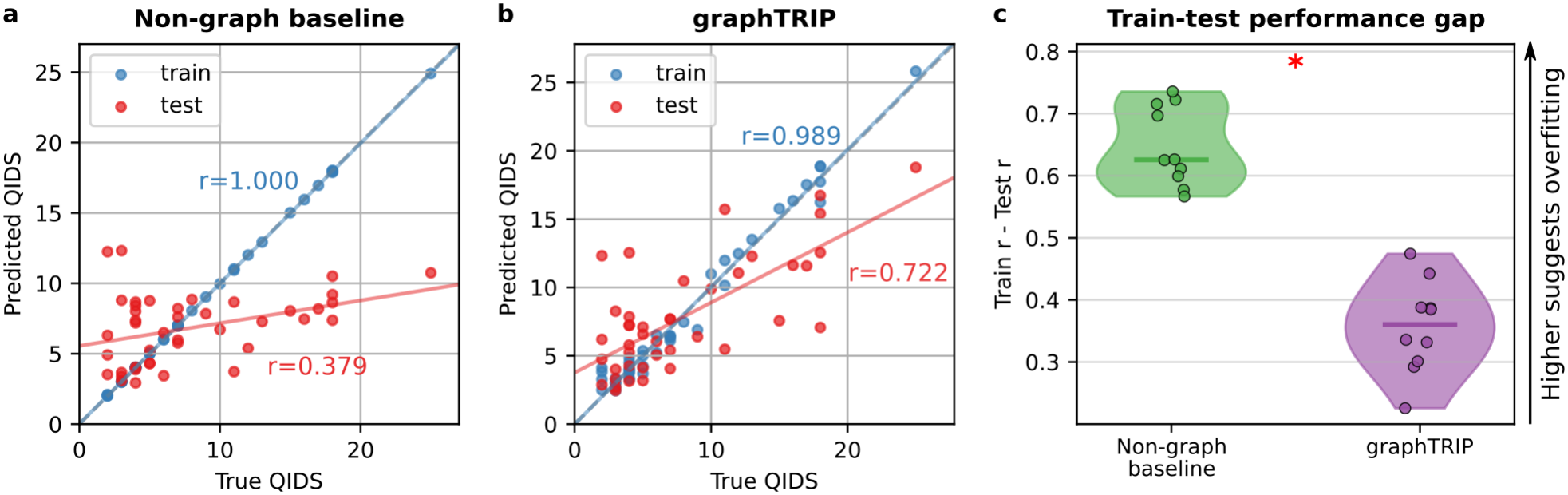
Weight sharing and regularisation mitigates overfitting relative to a non-graph autoencoder. Both models were trained with the same script, inputs, clinical covariates, loss weighting, and 7-fold cross-validation protocol. The key differences are with respect to architecture and regularisation. Architecture: `graphTRIP` uses a node-level VGAE that applies shared Graphormer parameters across brain regions (∼ 31k trainable parameters), so model capacity does not scale with the number of regions or connections. The non-graph baseline instead flattens each brain graph (i.e., concatenating the upper triangle of the FC adjacency with node features into a single vector) and encodes/decodes this vector with a standard multilayer autoencoder (3 hidden layers of width 512; ∼6.5M parameters), with no parameter sharing across regions or edges; its latent representations are graph-level vectors (not node-level as in `graphTRIP`), hence no pooling/readout over regions is used. Regularisation: `graphTRIP` is strongly regularised. It uses dropout (*p* = 0.25), L2 weight decay (*λ* = 0.01), and a KL regularisation term pulling the approximate posterior towards a standard normal prior, in both the encoder-decoder and the downstream regression head. The non-graph baseline is trained without any of these (dropout = 0, *λ* = 0, KL weight = 0, i.e. a deterministic-objective autoencoder) **a**, Non-graph baseline: perfect training fit (*r* = 1.00) but poor test generalisation (*r* = 0.38). **b**, `graphTRIP`: strong test performance (*r* = 0.72) and a smaller train-test gap. **c**, Across 10 random initialisations, the train-test gap (*r*_train_ − *r*_test_) is significantly smaller for `graphTRIP` and approximately halved relative to the non-graph baseline.

**FIG 16.**
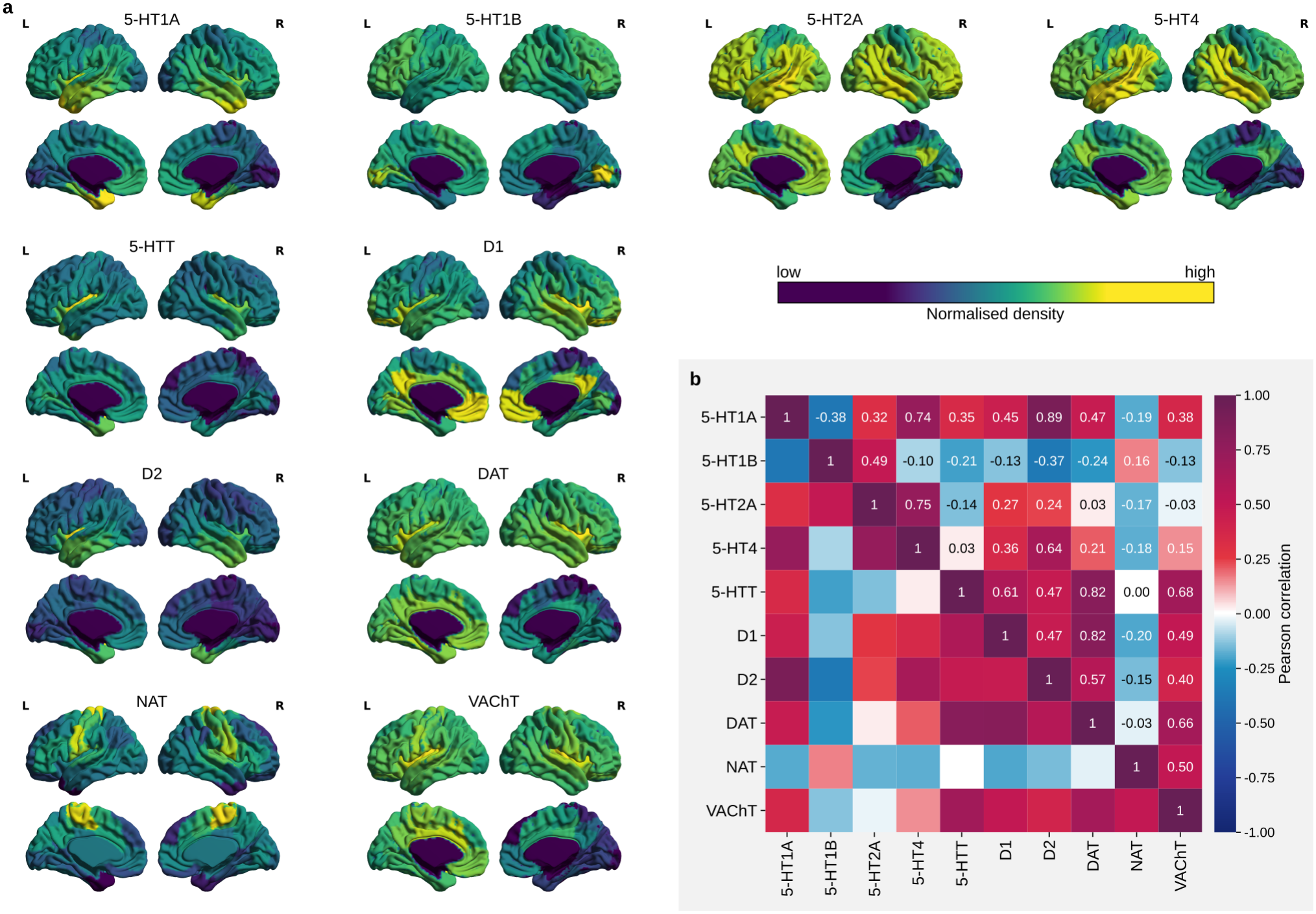
Normative molecular target distributions and cross-correlations. **a**, Normative molecular target maps used in this study. Each map is individually normalised; the colour scale therefore indicates relative density within a given target map and should not be compared across targets. **b**, Cross-correlation matrix showing spatial similarity between normative target maps. Each row and column represents a molecular target, and matrix entries indicate Pearson correlation coefficients between pairs of target distributions.

To formally test this relationship, we fitted an OLS regression model predicting regional attribution weights from each region’s position along the unimodal-transmodal axis together with ten normative molecular target density maps derived from open-access datasets [20, 41, 42] (see Methods). Model predictions correlated significantly with observed attribution patterns for both `graphTRIP` and `Medusa-graphTRIP` when assessed against 1,000 spatial autocorrelation–preserving null models [43] (`graphTRIP`: *r* = 0.4348, *p* = 0.0020; `Medusa-graphTRIP`: *r* = 0.8946, *p* = 0.0010). Dominance analysis identified the unimodal-transmodal axis as the strongest predictor in both cases, followed by dopamine, noradrenaline, and serotonin transporter densities for `graphTRIP` outcome predictions, and by serotonin receptor densities (5-HT2A, 5-HT4, and 5-HT1A) for `Medusa-graphTRIP` ITE estimates (Fig. 6a,b).

### Biomarkers of treatment responsiveness

Beyond regional attributions, we developed GRAIL to systematically link treatment response predictions to brain graph-derived biomarkers (Fig. 6c). This method exploits the dual structure of `graphTRIP` – a VGAE for learning brain graph representations and an MLP for predicting treatment outcome. For any differentiable biomarker of interest (*b*), GRAIL computes the alignment (cosine similarity) between the gradient of the model’s prediction (*ŷ*) and the gradient of *b* in latent space. Intuitively, these gradients describe how small changes in the latent brain representation affect the prediction *ŷ* and the biomarker *b*, respectively. Thus, their alignment provides a direct, analytical measure of learned associations: positive alignment indicates that higher biomarker values are associated with higher prediction values (e.g., higher post-treatment QIDS), and negative alignment suggests the opposite. Zero alignment suggests no learned association. Alignment significance was assessed by comparing observed values against null distributions generated using spatial autocorrelation–preserving spin permutations (see Methods).

We applied GRAIL to our models to assess learned associations for a range of candidate biomarkers, chosen to span two biologically and clinically relevant axes: large-scale brain networks (i.e., resting-state networks; RSNs [44]) and neuromodulatory systems. Specifically, we included ten normative molecular-target density maps available from open-access datasets [20, 42], covering major neurotransmitter systems implicated in antidepressant treatment [12]: serotonin (5-HT), dopamine (DA), noradrenaline (NA), acetylcholine (ACh). Candidate biomarkers included 1) mean brain-graph features (i.e., FC or REACT maps) within each RSN, and 2) spatial correlations between each brain graph feature and the normative molecular target maps. This broad screening strategy leverages GRAIL’s ability to discover biomarkers in a data-driven fashion.

To distinguish different types of biomarkers, we applied GRAIL in four complementary modes, corresponding to models trained to predict 1) overall response to either treatment (`graphTRIP`); 2) response to escitalopram (`Medusa-graphTRIP`’s escitalopram head); 3) response to psilocybin (`Medusa-graphTRIP`’s psilocybin head); and 4) ITEs (the difference between `Medusa-graphTRIP`’s heads). In the modes (1-3), gradient alignment quantifies whether a given biomarker is associated with better or worse response in general (1), or under the specific treatment (2-3). In mode (4), gradient alignment quantifies whether a biomarker is associated with relatively better response to escitalopram (positive alignment) or psilocybin (negative alignment). Importantly, by comparing the direction of alignment across these modes, biomarkers can be classified according to whether they predict treatment-invariant or drug-specific responsiveness or resistance. For example, a biomarker that showed significant negative alignment across modes (1-3), but no significant alignment with ITE predictions (mode 4), was classified as a shared responsiveness biomarker. Formal definitions of all biomarker categories are provided in the Methods.

We report in Fig. 6d only biomarkers with a valid category assignment in at least 50% of patients; all per-patient GRAIL category assignments are shown in Supp. Fig. 13. GRAIL identified multiple biomarkers associated with shared treatment responsiveness or resistance, as well as two biomarkers of differential response between psilocybin and escitalopram (Fig. 6d). Shared responsiveness biomarkers included higher network modularity within RSNs, and increased FC of regions with high normative 5-HT2A receptor density (FC corr 5-HT2A). In contrast, higher FC within VIS and SMN regions (FC mean VIS, FC mean SMN) was associated with overall treatment resistance. Interestingly, mean functional coupling between FPN regions and regions with high normative 5-HT1A and 5-HT2A receptor density (R5-HT1A mean FPN, R5-HT2A mean FPN) predicted resistance to psilocybin but response to escitalopram.

These biomarker findings are consistent with the input-domain ablations, in that treatment-specific REACT biomarkers of differential response were identified only by `Medusa-graphTRIP`.

Univariate biomarker-outcome correlations were consistent with the direction of GRAIL alignment (Supp. Fig. 14a). Across all 69 candidate biomarkers, group-averaged alignments correlated with biomarker-outcome correlations (*r* = 0.3896, *p* = 0.0009; Supp. Fig. 14b), whereas a linear ridge regression using all candidate biomarkers did not predict post-treatment QIDS (*r* = − 0.1995, *p* = 0.2052, *n* = 42). GRAIL profiles were highly reproducible among random-seed models trained on different train-test splits (`graphTRIP`: *r* = 0.9788; `Medusa-graphTRIP`: *r* = 0.9826), but distinct from those of 100 null models trained on shuffled labels (mean *r* = −0.0261 and −0.0737, respectively; Supp. Fig. 14c). In an out-of-fold analysis, training-set GRAIL alignments showed greater correspondence with held-out biomarker-outcome correlations for the identified biomarkers than for the remaining candidates (*p* = 0.0010, one-sided paired Wilcoxon test across seeds; Supp. Fig. 14d). Across all candidate biomarkers, `graphTRIP`’s training-set GRAIL alignments were significanlty sign-consistent with held-out biomarker-outcome associations (*p* = 0.0198, one-sided test against 100 label-permutation models; Supp. Fig. 14e).

## DISCUSSION

Predicting individual response to antidepressant treatment remains a major challenge in psychiatry, with implications for developing more effective, personalised interventions. Here, we introduced `graphTRIP`, a geometric deep learning (GDL) model that predicts post-treatment depression severity from pre-treatment neuroimaging and clinical data. Our model demonstrated strong prediction performance, generalised across brain parcellations, and maintained reconstruction accuracy in an independent cohort, with encouraging preliminary evidence of transferable outcome-predictive structure. Moreover, we developed `Medusa-graphTRIP`, a causal inference extension of `graphTRIP`, to estimate individual treatment effects under a counterfactual regression framework. Separately, we developed post-hoc interpretability analyses to identify biomarkers associated with these predictions.

`Medusa-graphTRIP` estimated an average treatment effect favouring psilocybin over escitalopram at three weeks post-treatment. This finding is consistent with previous analyses of the same dataset [33] and several other studies [10]. However, some patients were predicted to respond better to escitalopram. This underscores the importance of personalised treatment, and raises the question of what neurobiological mechanisms drive differential responsiveness to each drug.

To investigate this, we used three post-hoc inter-pretability analyses that address complementary questions at increasing specificity. Input-domain ablations quantify reliance on FC versus REACT. Regional attribution analysis identifies which brain regions most influence predictions via their latent representations (encoding both FC and REACT). Finally, GRAIL tests hypothesised candidate biomarkers (e.g. mean FC or REACT within a specific region or subnetwork) for directional alignment with predictions.

### Shared mechanisms of treatment responsiveness

Our analysis identified several brain-graph biomarkers that were associated with treatment outcome across both treatment arms, suggesting the presence of drug-invariant predictors of treatment responsiveness. Specifically, regional attribution analysis indicated that SMN and VIS regions contributed most strongly to `graphTRIP`’s outcome predictions. Consistent with this, GRAIL linked stronger mean FC in VIS and SMN regions to worse outcomes overall. This finding also aligns with recent work, reporting that baseline connectivity patterns involving VIS (and to a lesser extent SMN) networks are most informative for predicting early response to psilocybin [25].

One possible interpretation is that these biomarkers relate to individual differences in the brain’s functional cortical hierarchy. A prominent view posits that cortical organisation follows a unimodal-transmodal gradient, ranging from sensory systems to higher-order association cortices [38], and that psychedelic treatment may act in part by modifying or flattening this hierarchical organisation [45–47]. Beyond acute psychedelic effects on hierarchical organisation, baseline topography of the principal primary-to-transmodal connectome gradient has also been shown to predict symptomatic improvement under conventional antidepressant treatment [48]. In that light, our finding that stronger baseline coupling within unimodal systems such as VIS and SMN relates to poorer outcomes is consistent with the broader view that individual differences along the sensory-association axis are relevant to treatment responsiveness. Conversely, higher modularity and stronger coupling of regions associated with high normative 5-HT2A density – which tend to be located higher along the cortical hierarchy – may reflect a baseline organisation that supports better response to serotonergic interventions.

Together, these results suggest that baseline cortical hierarchy may contribute to treatment responsiveness in a manner that is partly shared across psilocybin and escitalopram. However, note that a drug-invariant baseline predictor does not imply that both treatments modify cortical hierarchy in the same way; indeed, recent evidence points to distinct hierarchical reconfigurations under psilocybin and escitalopram [27].

### Treatment-specific mechanisms of treatment responsiveness

Our analyses further suggested that treatment response heterogeneity is most strongly expressed in higher-order association systems. In particular, regional attribution analysis indicated that DMN and FPN regions contributed most to `Medusa-graphTRIP`’s ITE predictions. Additionally, GRAIL identified mean FPN REACT values for 5-HT1A and 5-HT2A as differential biomarkers, predicting better response to escitalopram but poorer response to psilocybin.

This pattern aligns with extensive literature implicating the FPN and DMN in depression and antide-pressant response [16, 47, 49, 50], and is also consistent with the view that psilocybin and SSRIs modulate serotonergic signalling through distinct mechanisms: Chronic SSRI use is thought to elevate cortical sero-tonin levels, which may bias signalling towards post-synaptic 5-HT1A-mediated inhibition and thereby reduce layer-V pyramidal cell excitability, particularly in higher-order association cortices where 5-HT1A is densely expressed [4, 51]. Such 5-HT1A-linked effects have been proposed to contribute to core therapeutic aspects of SSRI treatment, including improved stress regulation and emotional blunting [51]. In contrast, psilocybin acts as a direct agonist of excitatory 5-HT2A receptors, which are often co-expressed on the same layer-V pyramidal neurons that also express 5-HT1A [51]. A prominent hypothesis is that acute 5-HT2A activation transiently increases pyramidal cell excitability and initiates down-stream plasticity-related processes that support longer-lasting therapeutic effects [4].

Taken together, while the antidepressant mechanisms of both drugs remain incompletely understood, converging evidence suggests that they may modulate pyramidal cell excitability in association cortices via opposing 5-HT1A- and 5-HT2A-mediated effects. This aligns with our finding that FPN serotonergic coupling differentially predicts response to escitalopram versus psilocybin.

### Limitations and future work

A primary limitation of this study is the small sample size (*N* = 42). Given the high-dimensional brain graph inputs, it may seem surprising that a neural network can predict treatment outcomes from so few patients. One answer to this is that `graphTRIP` is a graph neural network (GNN). Rather than learning separate parameters for each brain region and connection, the model applies the same learned functions to every region and edge in each patient’s brain graph. This parameter sharing substantially reduces effective model complexity [52, 53] and is a central reason the approach is feasible in small neuroimaging cohorts. `graphTRIP` is also strongly regularised. A comparison to a matched non-graph autoen-coder baseline without weight sharing or regularisation shows that this design effectively mitigates overfitting (Supp. Fig. 15). Nevertheless, further evaluation in larger cohorts is needed to corroborate generalisability.

One potential source of confusion lies in the use of two separate models to predict treatment outcome (`graphTRIP`) and treatment effects (`Medusa-graphTRIP`). Conceptually, outcome prediction involves learning the two main effects of treatment and brain-graph features, whereas ITE estimation requires capturing the *interaction* between treatment and brain-graph features – that is, treatment-specific moderators. Our results suggest that `graphTRIP` captures the main effects well: after controlling for treatment, the correlation between true and predicted outcomes was slightly reduced but remained significant, indicating that both treatment and brain-graph effects were learned. However, `graphTRIP` likely captures treatment-specific interactions only partially. This can occur when a treatment-invariant predictive signal dominates the data, such that the model can achieve strong performance without learning additional treatment-specific features. In such settings, meta-learning theory suggests that models with separate predictors for each treatment arm can be better suited to capture heterogeneous treatment effects, albeit at the cost of higher variance when per-arm sample sizes are small [36]. `Medusa-graphTRIP` was therefore introduced to explicitly model treatment-specific outcome functions. Importantly, `graphTRIP` remains valuable in our setting because its single prediction head is trained on outcomes from both arms, providing a more stable estimator of overall treatment response and enabling the identification of drug-invariant biomarkers. Analysing inter-pretability results across both models allowed us to distinguish shared biomarkers from drug-specific moderators in a conservative manner – that is, supported by convergent findings across architectures with different inductive biases. With larger samples, this two-model strategy may no longer be necessary, and either `graphTRIP` or `Medusa-graphTRIP` alone may be sufficient.

Estimating causal treatment effects from observed data is inherently challenging because it requires reasoning about counterfactual outcomes which are never directly observed. Consequently, causal conclusions are only as reliable as the assumptions under which they are derived. One key assumption is ignorability, i.e., that treatment assignment is independent of potential outcomes conditional on baseline covariates [32]. While RCTs are designed to approximate this condition, small-sample effects can still introduce subtle imbalances. In our data, treatment assignment was modestly predictable from pretreatment variables, primarily due to slightly higher base-line QIDS in the psilocybin group (Supp. Fig. 11b,c). However, `Medusa-graphTRIP`’s ITE estimates were only weakly sensitive to baseline QIDS (Supp. Fig. 11d), suggesting that this imbalance is unlikely to be a major driver of our counterfactual predictions. Nevertheless, confirming treatment-specific biomarkers and improving the reliability of ITE estimation will require larger RCTs, ideally including comparisons across multiple antidepressant interventions. This would be a major step towards truly advancing personalised treatment of depression.

The GRAIL method also has certain constraints. First, it evaluates individual biomarkers, while treatment response likely depends on complex interactions. Second, GRAIL does not account for redundancy among correlated biomarkers; when two candidate biomarkers are highly correlated, both may show similar alignment patterns, but the method does not indicate which one is more explanatory. This limitation is particularly relevant given the substantial correlations among several normative molecular target maps used in our analysis (Supp. Fig. 16b). Furthermore, GRAIL identifies statistical relationships, not causal mechanisms. This distinction holds even when GRAIL is applied to `Medusa-graphTRIP`: while the model’s ITE estimates are causal (insofar as above-mentioned assumptions are met), GRAIL alignments remain associations between candidate biomarkers and the model’s learned predictions. As a result, correlated candidate biomarkers receive similar GRAIL scores even in a scenario where only one of them is causal. An exciting future direction involves integrating brain connectivity and molecular chemoarchitecture with generative whole-brain models – biophysically informed simulations of brain activity [54–59]. By using `graphTRIP` to predict treatment response from simulated brain dynamics and backpropagating through the whole-brain model, we could systematically screen for plausible neurophysiological mechanisms underlying treatment response.

Reconstruction of R5-HTT node features was consistently weaker than for FC, R5-HT1A and R5-HT2A (Fig. 2d). This is expected given the anatomy of the serotonin transporter, which is most densely expressed in midbrain, striatal and thalamic regions and less so in the neocortex, whereas 5-HT1A and 5-HT2A receptors show substantial cortical expression [42]. Since our primary analyses use a cortex-only Schaefer parcellation, reconstructing the R5-HT1A and R5-HT2A maps with stronger cortical structure is presumably easier for the VGAE than reconstructing R5-HTT.

Finally, we used normative molecular target maps derived from independent PET datasets, acquired in healthy individuals. While this is standard practice in molecular-enriched neuroimaging analyses [19], it is important to note that these maps were obtained from independent cohorts using distinct imaging methods. Although all data were transformed to standard space, such differences may introduce variability in resolution. Nonetheless, extensive work has demonstrated that this approach yields valid and clinically informative in-sights [18, 19, 22–24, 60, 61].

## CONCLUSION

In this work, we introduced `graphTRIP`, a geometric deep learning approach for predicting antidepressant treatment outcomes, offering both high predictive accuracy and interpretability. `graphTRIP` overcomes key limitations of conventional ML approaches, generalises across brain atlases, enables patient-specific biomarker discovery, and – via its causal extension – can estimate which treatment is likely to work best for each patient. Our analysis highlights the role of unimodal brain regions in early treatment response to both escitalopram and psilocybin, and reveals serotonergic signalling in association networks as a treatment-specific moderator. Future work could scale our approach to larger datasets and integrate whole-brain models to move beyond biomarker-based to mechanistic explanations of treatment responsiveness. Overall, these advances mark a step toward data-driven, personalised treatment selection, bringing ML models closer to clinical translation.

## METHODS

### Datasets

We used data from two clinical trials conducted at the Imperial Clinical Research Facility and approved by relevant UK regulatory bodies. All participants provided written informed consent. The main dataset consisted of a double-blind randomised controlled trial (DB-RCT, clinicaltrials.gov: NCT03429075) comparing psilocybin and escitalopram for the treatment of major depressive disorder (MDD). An independent dataset was derived from an open-label trial of psilocybin treatment (gtr.ukri.org: MR/J00460X/1). Detailed trial protocols and clinical outcomes have been previously published [47].

### Participants

Eligible participants in both trials were diagnosed with unipolar MDD (Hamilton Depression Rating Scale score ≥ 16). The DB-RCT included 59 patients, randomly assigned to either psilocybin (N = 30) or escitalopram (N = 29) treatment. The final imaging sample comprised 22 patients in the psilocybin arm (mean age = 41.9*±* 11.0 SD years; 8 female) and 20 in the escitalopram arm (mean age = 38.7 11.0 SD years; 6 female). The open-label trial included 19 patients with treatment-resistant depression (TRD), defined as having failed to respond to multiple courses of antidepressant treatment (mean = 4.6 *±* 2.6 SD past medications). Out of the initial sample, 16 patients were retained for analysis after excluding three due to excessive head motion (mean age = 42.75 *±* 10.15 SD years; 4 female). Exclusion criteria for both trials included a history of psychosis, significant medical conditions, serious suicide attempts, pregnancy, and MRI contraindications. The DB-RCT additionally excluded patients with contraindications for SSRIs or prior escitalopram use.

### Treatments

In both trials, all participants underwent a pretreatment baseline session, involving clinical assessment and resting-state fMRI. The psilocybin arm received a 25 mg dose on Day 1 followed by a second identical dose 3 weeks later, along with daily placebo capsules for six weeks. The escitalopram arm received a negligible 1 mg psilocybin dose on Day 1, followed by 10 mg of escitalopram daily for the first 3 weeks, increased to 20 mg daily thereafter. In the open-label trial, patients received two psilocybin doses (10 mg and 25 mg, one week apart).

### Clinical outcome measures

Depression severity was assessed using both the Quick Inventory of Depressive Symptomatology (QIDS) and the Beck Depression Inventory (BDI-1A) in both trials, at baseline and post-treatment. In the DB-RCT, QIDS was the primary outcome measure, with post-treatment assessments conducted 3 weeks after dosing day 2. BDI served as a secondary outcome measure in this trial. Thus, we used QIDS as the default prediction target of our model. The open-label trial used BDI as the primary outcome measure, with post-treatment assessments occurring 1 week after dosing day 2. In the open-label dataset, additional pre-treatment scores from the HAMD and LOT-R scales were available, which we included as input features when training models (simple MLP or `graphTRIP`) from scratch without pretraining.

### fMRI data acquisition and preprocessing

Resting-state fMRI data were acquired using a 3T Siemens Tim Trio scanner with T2*-weighted echo-planar imaging. In the DB-RCT, scans included 480 volumes in approximately 10 min (TR = 1,250 ms; TE = 30 ms; 44 axial slices; spatial resolution = 3 mm isotropic; flip angle = 70 degrees; bandwidth = 2,232 Hz per pixel; and GRAPPA acceleration = 2). In the open-label trial, scans included 280 volumes in approximately 8 min (TR = 2,000 ms; TE = 31 ms; 36 axial slices; flip angle = 80 degrees; bandwidth = 2,298 Hz per pixel; and GRAPPA acceleration = 2). We used the Schaefer et al. [34] brain atlas with 100 parcels as the default atlas, and also conducted analyses using the Schaefer atlas with 200 parcels and the Automated Anatomical Labeling (AAL) atlas [35]. Preprocessing was performed using a custom pipeline integrating FSL, AFNI, Freesurfer, and ANTs, following standard procedures: motion correction, spatial smoothing, band-pass filtering (0.01–0.08 Hz), and nuisance regression. Volumes with *>* 20% framewise displacement *>* 0.5 mm were excluded. Full details of the preprocessing steps can be found in [47].

### Functional connectivity

BOLD time series were parcellated and z-scored using the Schaefer atlases with 100 or 200 parcels [34], and the AAL atlas [35]. Functional connectivity (FC) was then computed as the Pearson correlation coefficient between the z-scored time series of each pair of brain regions, using Nilearn’s `ConnectivityMeasure` function (`kind=‘correlation’`). This resulted in an *N × N* FC matrix for each participant and parcellation.

### Molecular target maps

Positron emission tomography (PET) maps of regional neuroreceptor and transporter densities were obtained from publicly available normative datasets acquired in healthy individuals; no PET imaging was performed on trial participants. Serotonin receptor (5-HT1A, 5-HT2A) and serotonin transporter (5-HTT) maps were downloaded from Beliveau et al. [42] (https://nru.dk/index.php/allcategories/category/90-nru-serotonin-atlas-and-clustering). The maps were resampled to 2 mm MNI152 space, cerebellar voxels were excluded, and density values were normalised to a range of 0–1.

Additionally, we used PET maps for seven other molecular targets to explain the regional attention weight patterns and compute interpretable biomarkers in the gradient alignment analysis. Specifically, we included density maps for serotonin receptors (5-HT4, 5-HT1B), dopamine receptors (D1, D2) and transporter (DAT), noradrenaline transporter (NAT), and vesicular acetylcholine transporter (VAChT). These data were obtained from Hansen et al. [20].

### REACT maps

The REACT maps, which served as the node attributes of our main model, were computed for each participant using the voxelwise PET maps of 5-HT1A, 5-HT2A, and 5-HTT densities from Beliveau et al. [42]. The analysis was performed with the `react-fmri` Python toolbox [18], following the protocol described in Lawn et al. [19].

Specifically, we used the `react masks` command to generate masks that ensured that all voxels included in the subsequent analysis had valid PET data for all molecular targets, BOLD data for all participants, and were located within grey matter. Subsequently, we computed the REACT maps with the `react` command, which involves two sequential linear regressions for each voxel and molecular target. First, a voxel-level spatial regression of each molecular density map (5-HT1A, 5-HT2A, and 5-HTT) against the BOLD values at each time point yields a time series capturing the dominant fluctuations within each molecular system over time. In the second step, these molecular time series are regressed against the BOLD time series of each voxel, producing estimates of the coupling between each voxel’s activity and the broader activity associated with each molecular system. The resulting voxelwise REACT maps were parcellated to obtain one value for each brain region, participant, and atlas.

### Statistical analysis

Statistical analyses were performed in Python (version 3.12). Prediction accuracy was quantified as the Pearson correlation (*r*), *R*^2^, mean absolute error (MAE) and root-mean-square error (RMSE) between true and predicted outcomes. Significance of `graphTRIP` and `Medusa-graphTRIP` prediction performance was assessed against an empirical null distribution by retraining the entire model on shuffled post-treatment QIDS labels. We trained on 100 different label permutations, and for each permutation we trained 10 random seed models, where the seed determines train-test split and weight initialization. This matches how the main reported metrics are constructed (as mean test predictions over 10 seeds). Rank *p*-values were computed as *p* = (1 + # {null ≥ observed })*/*(1 + 100), so the lowest achievable *p*-value is *p <* 0.01. Unless stated otherwise, p-values are two-sided. Where indicated, p-values were corrected for multiple comparisons using false discovery rate (FDR) correction.

Model comparisons used paired Wilcoxon signed-rank tests across matched random seeds, which share the same train-test partitions. Furthermore, where the number of predictors is small compared to the sample size, we used ordinary least squares (OLS) regression; otherwise we used ridge regression. In the dominance analysis of regional attribution values, OLS was used to quantify explained variance across brain regions.

### Model and training configurations

Full hyperparameter configurations of all models, including architectural details and training parameters are provided in Supplementary Tables III-IV. Additionally, the complete codebase and all configuration files are available at: https://github.com/Imperial-MIND-lab/graphTRIP. Pseudo code, describing the `graphTRIP` training pipeline is provided in the Supplementary 27.

### graphTRIP model

The main model in our study, called `graphTRIP` model, consists of a variational graph autoencoder (VGAE) with graph attention layers and a downstream multi-layer perceptron (MLP). The VGAE is trained to learn informative latent node representations by reconstructing the original input graph, while the MLP predicts post-treatment QIDS scores based on these latent representations, the drug condition, and additional clinical data.

#### VGAE inputs

The input to the VGAE is a brain graph, where nodes represent brain regions defined by a brain atlas and edges represent FC between regions. Each node has a 3-dimensional feature vector, encoding the REACT values for 5-HT1A, 5-HT2A, and 5-HTT. Additionally, three conditional node features encode the 3D spatial coordinates of each brain region in MNI space. These conditional features provide the model with anatomical information about regional identity, while preserving permutation invariance of the graph representation. Since these features are normative and do not contain patient-specific information, they are used only by the encoder and are not reconstructed by the decoder. Edges exist between regions if the absolute FC value is among the global top 20% strongest connections to obtain a fixed connection density of *ρ* = 0.2 for each brain graph, and edge attributes are the original (non-thresholded) FC values. This thresholding approach was shown to yield robust and reliable network topology [31]. Self-connections were set to 1.

#### VGAE architecture

The VGAE consists of a node-level encoder and decoder and a graph-level readout layer. We will describe each component in turn.

##### Encoder

The encoder is based on a Graphormer-style architecture [62], which extends the transformer architecture to graph-structured data. Each node’s feature vector *x*_*i*_ is first concatenated with its conditional node features *c*_*i*_ (the MNI coordinates) and projected into a shared embedding space as

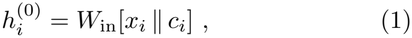

where *W*_in_ is a learnable linear projection.

The model then applies three Graphormer layers.At each layer *l*, node embeddings are updated using multi-head self-attention with two learned attention biases, derived from shortest-path distances between nodes and edge attributes (the FC weights). Specifically, the shortest-path bias 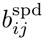 is obtained by embedding thediscretised shortest-path distance between nodes *i* and *j*, while the edge bias 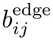 is computed by passing the corresponding FC value through a two-layer MLP with ReLU hidden-layer activation. Both biases are added to the attention scores *e*_*ij*_ between nodes *i* and *j* in a head-specific manner as follows.

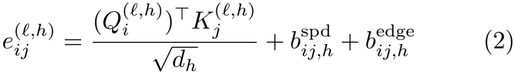

Here, *h* indexes attention heads, 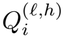 and 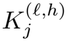 arelinear projections of the node embeddings, and *d*_*h*_ is the head dimension.

Attention weights are obtained via softmax normalisation over all nodes within the same graph,

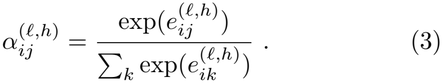

After applying a dropout layer, the normalised scores are used to compute the attention output as

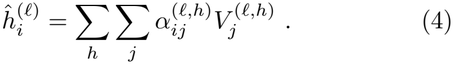

where 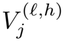 is another linear projection of the node embeddings.

The outputs from all attention heads are then linearly projected to allow mixing of information across heads,

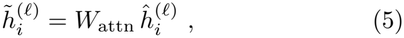

where *W*_attn_ is a learnable projection matrix.

Each Graphormer layer further includes a position-wise feedforward network with residual connections, layer normalisation, and dropout. Concretely, the layer update takes the form

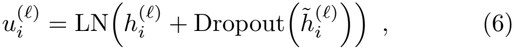

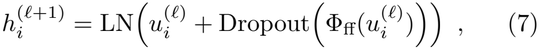

where LN( ·) denotes layer normalisation and Φ_ff_(·) is a two-layer MLP with ReLU activation in the hidden layer.

After the final Graphormer layer, node embeddings are projected to a lower-dimensional space,

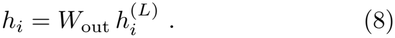

A final linear layer maps each node embedding *h*_*i*_ to a Gaussian distribution parameterised by a mean *µ*_*i*_ and log-variance log 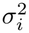,

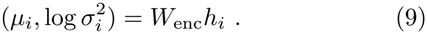

Latent node representations *z*_*i*_ are then sampled using the reparameterisation trick,

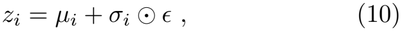

where *ϵ* ∼ *N* (0, *I*) and *σ*_*i*_ = exp(0.5 *·*log *σ*^2^). These latent samples serve as input to the decoders and the graph-level readout.

##### Decoder

The VGAE decoder consists of three independent sub-modules:

- **Node decoder:** A 3-layer MLP with LeakyReLU activation and dropout, and a linear output layer to reconstruct node features from *z*_*i*_.
- **Edge index decoder:** A 3-layer MLP with LeakyReLU activation and dropout, and a sigmoid output to predict the probability of edges between nodes.
- **Edge attribute decoder:** A 3-layer MLP with LeakyReLU activation and dropout, and a tanh output to predict FC values in the range [−1, 1].

The reconstruction of the thresholded FC matrix is obtained by elementwise multiplication of the edge-index and edge-attribute decoder outputs.

##### Readout layer

The VGAE readout layer aggregates the latent node vectors {*z*_*i*_*}* into graph-level representa-tions. Before aggregating, it first applies a single multihead self-attention block to the latent node vectors to allow each node representation to integrate information from other regions. This block works analogously to the encoder attention mechanism described above, but without the structural attention biases, and – again, just like in the encoder – it is followed by a position-wise feedfor-ward network (a two-layer MLP) with residual connections, layer normalisation, and dropout.

The resulting contextualised node embeddings are then aggregated using mean pooling to obtain a single graphlevel representation,

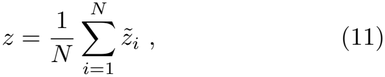

where 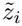 denotes the contextualised latent embedding of node *i* and *N* is the number of nodes in the graph.

#### Atlas-agnostic prediction

Graph neural networks are not tied to a fixed number of nodes. In `graphTRIP`, the encoder and decoder apply the same learned functions to every node and edge, so the number of trainable parameters does not depend on the number of parcels *N* . Across atlases, the numbers of nodes (*N* ) and edges change, but the per-node feature dimension remains fixed. So the encoder produces a fixed-size representation for each node by applying the same function *N* times. Mean pooling then aggregates these *N* latent node vectors into a fixed-size graph representation *z*, which serves as the input to the MLP prediction head. Because *z* is invariant to *N*, the prediction head does not need resizing or retraining when the atlas changes.

For this transfer to be meaningful in brain networks, nodes must encode anatomical location; otherwise shared node functions would treat brain regions as fully exchangeable. We therefore include each region’s MNI coordinates as conditional node features (see VGAE inputs), allowing the network to learn to distinguish prefrontal association cortices from occipital visual regions, for example.

#### MLP prediction head

The input to the MLP prediction head consists of a concatenation of the graph representation vector *z* and clinical data: baseline BDI and QIDS scores, a binary indicator of prior SSRI use (0 for no, 1 for yes), and the drug condition *d* (-1 for escitalopram, 1 for psilocybin). The MLP has three layers with LeakyReLU activation and dropout in the hidden layers (see Tab. III), and a linear output layer to predict post-treatment QIDS scores.

### Medusa-graphTRIP model

To estimate individual treatment effects (ITEs), we developed `Medusa-graphTRIP`, a causal extension of `graphTRIP` with separate prediction heads for each treatment condition. The model is inspired by the counterfactual regression framework introduced by Shalit et al. [32].

`Medusa-graphTRIP` employs a single VGAE to learn a shared latent brain-graph representation across patients from both treatment groups. The VGAE architecture is identical to that used in `graphTRIP`, with the exception of the readout layer (Tab. IV). Specifically, `Medusa-graphTRIP` uses a simple mean-standard deviation pooling, where the per-graph representation is formed by concatenating the mean and standard deviation of latent node vectors. We chose this less expressive pooling module to ensure that the two treatment-specific prediction heads receive statistically comparable latent representations.

Downstream of the shared VGAE, `Medusa-graphTRIP` includes two MLP prediction heads, each trained exclusively on patients from a single treatment condition. As in `graphTRIP`, the MLP heads receive the latent brain-graph representation together with baseline clinical data (baseline QIDS, baseline BDI, and SSRI history) and are trained to predict post-treatment QIDS scores. Treatment condition is not provided as an explicit input to the MLPs, but is used solely to route each patient’s latent representation to the appropriate treatment-specific head during training.

Following Shalit et al. [32], individual treatment effects for each patient *i* were estimated as the difference between the predictions of the two heads,

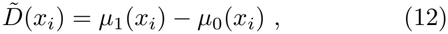

where 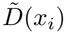 denotes the estimated ITE, *x*_*i*_ represents the patient’s covariates, and *µ*_0_ and *µ*_1_ are the prediction heads trained on escitalopram and psilocybin patients, respectively. Thus, *µ*_0_ estimates response under escitalopram, while *µ*_1_ estimates response under psilocybin.

### Model training

Unless stated otherwise, `graphTRIP` and `Medusa-graphTRIP` were trained using identical training protocols (pseudocode in Supp. 27). All models were implemented in PyTorch and trained using the Adam optimiser with a learning rate of 0.001.

#### Training objective

The VGAE and downstream MLP(s) were trained jointly to optimise both graph reconstruction and outcome prediction. Because training was performed end-to-end, gradients from the outcome-prediction loss propagated through the MLP into the VGAE encoder, thereby shaping the learned latent representation. For `graphTRIP`, the total training objective was defined as a weighted combination of the VGAE loss and the MLP loss,

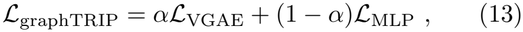

with *α* = 0.5.

For `Medusa-graphTRIP`, an additional maximum mean discrepancy (MMD) penalty was used to encourage similar latent representations across treatment groups, as recommended by Shalit et al. [32]:

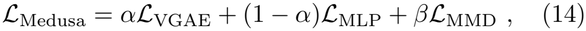

where *β* = 1 controls the strength of the MMD penalty. The MMD was computed between latent representations of patients receiving psilocybin and escitalopram using a Gaussian radial basis function kernel with bandwidth estimated from the data.

The VGAE loss comprises a reconstruction term, a Kullback-Leibler (KL) divergence term, and an L2 regularisation term on the VGAE parameters,

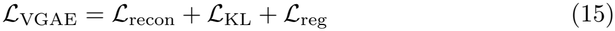

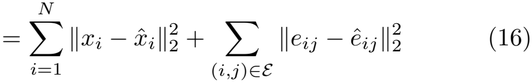

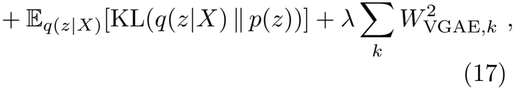

where *x*_*i*_ and 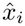 denote the original and reconstructed node features, *e*_*ij*_ and *ê*_*ij*_ the original and reconstructed edge attributes (FC values), *N* is the number of nodes, and denotes the set of unique (upper-triangular) edges. The final term corresponds to standard weight-decay regularisation over all VGAE parameters *W*_VGAE_ with *λ* = 0.01.

The MLP loss consists of a mean squared error prediction term and an L2 weight-decay regularisation term,

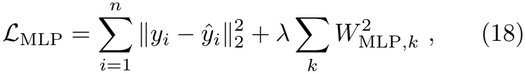

where *y*_*i*_ and *ŷ*_*i*_ are the true and predicted post-treatment QIDS scores for the *i*-th patient in a batch of size *n, λ* = 0.01 is the regularisation strength, and *W*_MLP_ denotes the MLP parameters.

#### Training protocol

All models were trained for 300 epochs using 7-fold cross-validation (CV), with a batch size of 9. CV splits were stratified by treatment condition to maintain balanced proportions of psilocybin and escitalopram patients in each fold.

Latent-sample augmentation was used only for `Medusa-graphTRIP`. Specifically, during training, the MLP prediction heads received three independently sampled latent representations per patient. The rationale behind this was to slightly augment the effective training data for the treatment-specific MLP heads, which are trained on fewer samples compared to `graphTRIP`’s single MLP head due to treatment stratification.

QIDS predictions and reconstructions from a given model were always generated only for patients that the model did not train on. For within-cohort evaluations this means test-fold patients; for zero-shot cross-cohort evaluations, all patients in the independent dataset were never seen by any model. The only exception are the interpretability analyses (regional attributions, GRAIL), where we interpreted the predictions of each model for all patients (train and test set). Each full training procedure was repeated ten times with different random seeds. Unless stated otherwise, all reported predictions correspond to mean test-set predictions averaged across these ten runs.

### Baseline-severity input normalization for the independent cohort

For zero-shot outcome prediction on the independent TRD cohort, drug condition and prior SSRI use were set to 1 for all patients (psilocybin-only, treatment-resistant). Baseline QIDS and BDI were affine-mapped onto each fold model’s expected input scale before MLP input; brain-graph inputs and model parameters remained fixed. Raw baseline scores *x* were transformed as

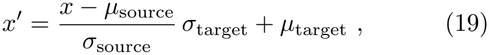

where (*µ*_target_, *σ*_target_) are the mean and standard deviation of the attribute on the training subjects of fold *k*, and (*µ*_source_, *σ*_source_) are leave-one-out statistics computed from the other patients in the independent cohort, so no target patient contributed to its own renormalization. This unsupervised approach is equivalent to adaptive batch normalization [63], applied only at the input layer, only for baseline severity inputs, and with leave-one-out statistics. The same protocol was applied to the clinical-only MLP baseline.

### Gradient Alignment for Interpreting Latent-variable Models (GRAIL)

The GRAIL method was developed to estimate learned associations between candidate brain-graph-derived biomarkers and model predictions (e.g. post-treatment QIDS or ITEs). Positive alignment indicates that higher values of a biomarker are associated with a higher prediction value, while negative alignment indicates the opposite. An alignment of zero indicates no association between the biomarker and predicted treatment response.

GRAIL leverages the two-part architecture of the `graphTRIP` and `Medusa-graphTRIP` models, consisting of a VGAE and an MLP prediction head. The VGAE encoder maps input brain graphs to latent representations *z*, which serve as inputs to both the VGAE decoder (reconstructing brain graphs) and the MLP (predicting post-treatment QIDS or ITEs). Biomarkers of interest (e.g., mean FC within an RSN) are computed from recon-structed graphs. The only requirement for a biomarker is that it can be expressed as a differentiable function of the reconstructed brain graph.

GRAIL computes the latent gradients of the MLP prediction *ŷ* and the candidate biomarker *b*, i.e., ∇ _*z*_*ŷ* and the ∇ _*z*_*b*. These gradients describe directions in latent space that locally increase the predicted outcome and the biomarker value, respectively. Alignment between prediction and biomarker is quantified as the cosine similarity between the gradients,

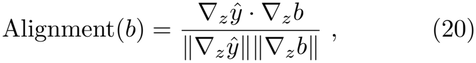

which ranges from −1 (maximal negative alignment) to 1 (maximal positive alignment).

A first-stage screening ensured that only biomarkers with stable local associations around the patient’s latent representation were considered: For each patient and trained model, we generated 25 samples from a Gaussian distribution centred at the patient-specific latent mean. For each latent sample, we computed the predicted outcome (or ITE), all candidate biomarkers, their respective gradients, and the corresponding alignment values. For each biomarker, we then performed a one-sample *t*-test across the 25 latent samples to test whether alignment values were significantly different from zero. Biomarkers with a Cohen’s *d >* 0.5 and FDR-corrected *p <* 0.05 were retained for further analysis. We stored the mean alignment across latent samples for all biomarkers.

For biomarkers passing the first-stage screening, statistical significance was assessed using spatially informed null models. All candidate biomarkers were derived from either regional assignments to RSNs, or normative molecular target density maps. Accordingly, surrogate biomarkers were generated, for each latent sample, using 1000 spatial-autocorrelation preserving spin permutations of RSN assignments or normative density maps. Gradient alignments were computed for each sur-rogate biomarker in the same manner as for the observed biomarker and then averaged across latent samples. This yielded a null distribution of mean alignment values for each candidate biomarker. Observed mean alignment values were compared against these null distributions, and biomarkers were deemed significant if their alignment exceeded the null distribution after FDR correction at *p <* 0.05.

The above procedure was applied independently to all models trained with different random seeds and CV folds. For each patient and biomarker, this resulted in a binary indicator of whether the biomarker was found to be significantly aligned in a given model. To identify biomarkers that were robustly associated with predictions across model initializations, we tested whether the number of significant detections exceeded chance expectation. Specifically, for each patient and biomarker, we computed the probability of observing *k* or more significant detections out of *n* trained models under a binomial model,

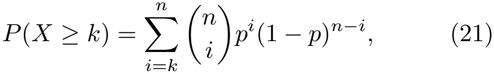

where *p* denotes the nominal false-positive rate. Biomarkers with low tail probabilities were interpreted as robustly aligned across random initializations for a particular patient. For reporting alignment values, we computed performance-weighted mean alignments across models, with weights proportional to test-fold prediction accuracy (true-vs-predicted Spearman correlation).

We performed four complementary checks of GRAIL associations:

1. We compared group-mean alignments with univariate biomarker-outcome correlations across candidate biomarkers (Supp. Fig. 14a,b).
2. We assessed profile reproducibility by splitting the ten random-seed `graphTRIP` or `Medusa-graphTRIP` models into all possible groups of five, and correlating their patient-by-biomarker GRAIL alignment profiles; as a negative control, we then compared `graphTRIP`’s alignment profile with those from the 100 null models trained on permuted labels (Supp. Fig. 14c).
3. For each CV-fold `graphTRIP` model with positive test-fold prediction performance, we averaged GRAIL alignments over its training-set patients and correlated this vector with biomarker-outcome correlations, calculated only in its held-out test-set patients. We performed this analysis once including only the identified biomarkers, and once including the remaining candidates. This yields two correlation values per CV-fold model. We then averaged those across CV-fold models within each seed, and compared mean correlations of identified versus remaining candidates (Supp. Fig. 14d).
4. Without selecting any biomarkers, we quantified the sign consistency between training-set GRAIL alignments and test-set biomarker-outcome correlations across all candidate biomarkers. For each CV-fold model: 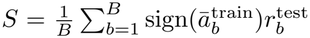, where *B* = 69, 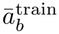 is the mean training-set GRAIL alignment for biomarker *b*, and 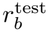 isits biomarker–outcome correlation in the held-out patients. We compared the mean *S* across `graphTRIP` CV-fold models with the corresponding means from 100 label-permutation null models (Supp. Fig. 14e).

### Biomarker categories

To distinguish shared versus drug-specific biomarkers of treatment responsiveness, we applied GRAIL in four complementary modes: 1) `graphTRIP` (shared outcome prediction across both treatment arms); 2) the psilocybin head of `Medusa-graphTRIP`; 3) the escitalopram head of `Medusa-graphTRIP`; and 4) `Medusa-graphTRIP`’s ITE predictions. For outcome-prediction models (1-3), positive alignment indicates biomarkers associated with higher predicted post-treatment QIDS (treatment resistance), whereas negative alignment indicates biomarkers associated with lower predicted post-treatment QIDS (treatment responsiveness). For the ITE mode (4), positive alignment indicates biomarkers associated with relatively better predicted response to escitalopram, whereas negative alignment indicates relatively better predicted response to psilocybin.

For each patient, we assigned each biomarker to a category based on the combination of significant alignment directions across the four modes (Table I). Biomarkers whose alignment patterns did not match any valid combination were considered invalid and remained unassigned. For downstream analyses, we retained biomarkers that were assigned a valid category in at least 50% of patients.

**TABLE 1.** Valid biomarker category assignments based on GRAIL alignment direction across four modes. ‘–’ indicates significant negative alignment, ‘+’ indicates significant positive alignment, and ‘0’ indicates non-significant alignment.

| Shared | Psilocybin | Escitalopram | ITE | Biomarker category |
|---|---|---|---|---|
| – | – | – | 0 | Shared responsiveness |
| – | – | – | + | Shared responsiveness |
| – | – | – | – | Shared responsiveness |
| + | + | + | 0 | Shared resistance |
| + | + | + | + | Shared resistance |
| + | + | + | – | Shared resistance |
| – | 0 | – | + | Escitalopram responsiveness |
| 0 | 0 | – | + | Escitalopram responsiveness |
| – | – | 0 | – | Psilocybin responsiveness |
| 0 | – | 0 | – | Psilocybin responsiveness |
| + | 0 | + | – | Escitalopram resistance |
| 0 | 0 | + | – | Escitalopram resistance |
| + | + | 0 | + | Psilocybin resistance |
| 0 | + | 0 | + | Psilocybin resistance |
| 0 | + | – | + | Escitalopram responsiveness & psilocybin resistance |
| 0 | – | + | – | Psilocybin responsiveness & escitalopram resistance |

**TABLE 2.**
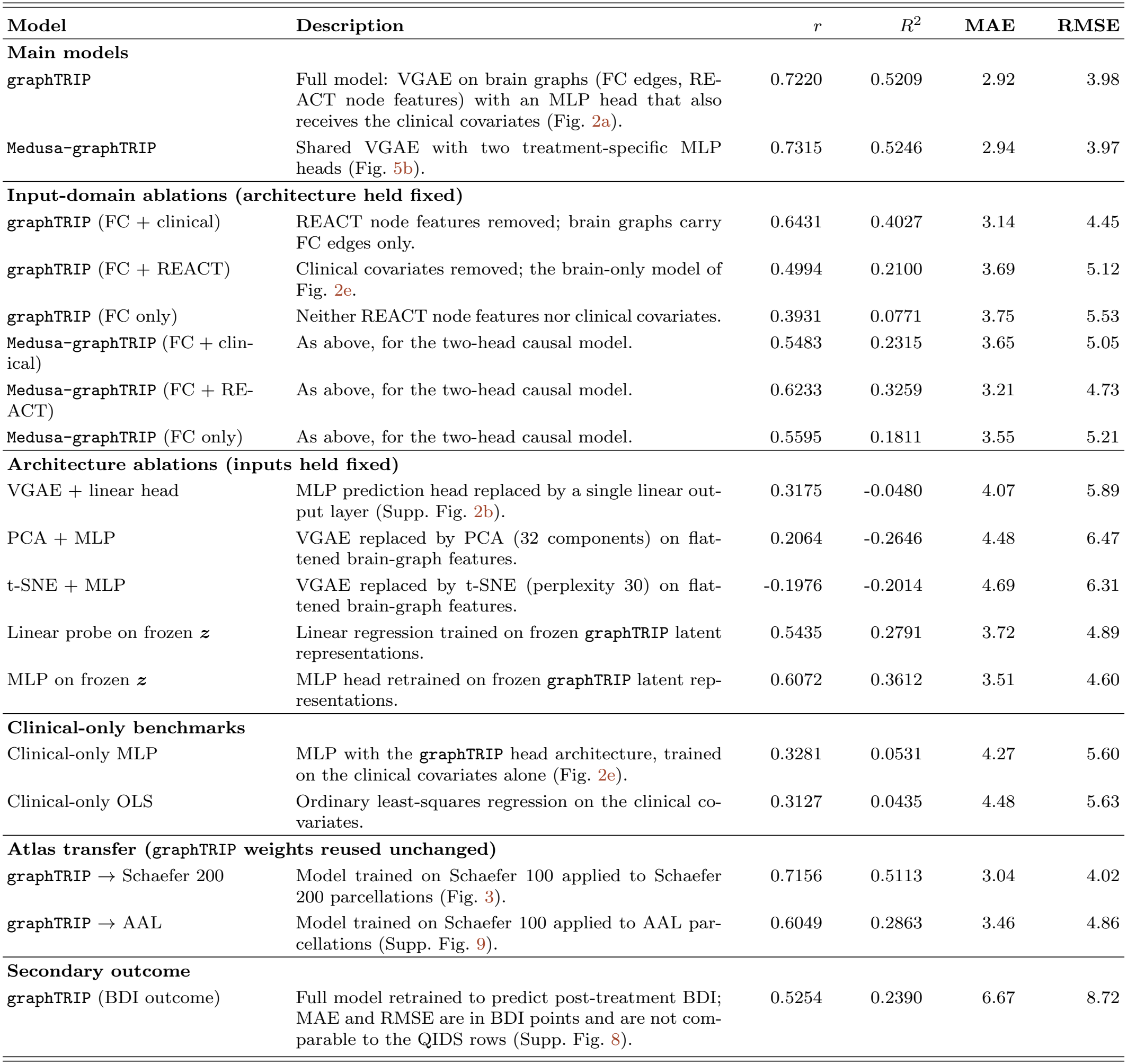
Prediction performance of all models. All models were evaluated on the primary dataset (*N* = 42) using the same 7-fold cross-validation protocol, and all metrics were computed from test-fold predictions averaged across 10 random seeds. Clinical covariates denotes baseline QIDS, baseline BDI and SSRI history; all models additionally receive the drug condition as an input. *r*: Pearson correlation between true and predicted outcomes. *R*^2^ = 1 −SSE*/*SST is computed against the sample mean, so that negative values indicate predictions worse than always predicting the sample mean. MAE and RMSE are reported in QIDS points (scale range 0–27), except where stated otherwise. Inferential *p*-values for prediction performance are reported in the main text for `graphTRIP`, `Medusa-graphTRIP`, and `graphTRIP` atlas transfers, where they are obtained from label-permutation nulls (Methods); other models are reported descriptively.

| **Model** | **Description** | *r* | *R*<sup>2</sup> | **MAE** | **RMSE** |
|---|---|---|---|---|---|
| **Main models** |  |  |  |  |  |
| graphTRIP | Full model: VGAE on brain graphs (FC edges, RE-ACT node features) with an MLP head that also receives the clinical covariates (Fig. 2a). | 0.7220 | 0.5209 | 2.92 | 3.98 |
| Medusa-graphTRIP | Shared VGAE with two treatment-specific MLP heads (Fig. 5b). | 0.7315 | 0.5246 | 2.94 | 3.97 |
| **Input-domain ablations (architecture held fixed)** |  |  |  |  |  |
| graphTRIP (FC + clinical) | REACT node features removed; brain graphs carry FC edges only. | 0.6431 | 0.4027 | 3.14 | 4.45 |
| graphTRIP (FC + REACT) | Clinical covariates removed; the brain-only model of Fig. 2e. | 0.4994 | 0.2100 | 3.69 | 5.12 |
| graphTRIP (FC only) | Neither REACT node features nor clinical covariates. | 0.3931 | 0.0771 | 3.75 | 5.53 |
| Medusa-graphTRIP (FC + clin-ical) | As above, for the two-head causal model. | 0.5483 | 0.2315 | 3.65 | 5.05 |
| Medusa-graphTRIP (FC + RE-ACT) | As above, for the two-head causal model. | 0.6233 | 0.3259 | 3.21 | 4.73 |
| Medusa-graphTRIP (FC only) | As above, for the two-head causal model. | 0.5595 | 0.1811 | 3.55 | 5.21 |
| **Architecture ablations (inputs held fixed)** |  |  |  |  |  |
| VGAE + linear head | MLP prediction head replaced by a single linear out-put layer (Supp. Fig. 2b). | 0.3175 | -0.0480 | 4.07 | 5.89 |
| PCA + MLP | VGAE replaced by PCA (32 components) on flat-tened brain-graph features. | 0.2064 | -0.2646 | 4.48 | 6.47 |
| t-SNE + MLP | VGAE replaced by t-SNE (perplexity 30) on flat-tened brain-graph features. | -0.1976 | -0.2014 | 4.69 | 6.31 |
| Linear probe on frozen ***z*** | Linear regression trained on frozen graphTRIP latent representations. | 0.5435 | 0.2791 | 3.72 | 4.89 |
| MLP on frozen ***z*** | MLP head retrained on frozen graphTRIP latent rep-resentations. | 0.6072 | 0.3612 | 3.51 | 4.60 |
| **Clinical-only benchmarks** |  |  |  |  |  |
| Clinical-only MLP | MLP with the graphTRIP head architecture, trained on the clinical covariates alone (Fig. 2e). | 0.3281 | 0.0531 | 4.27 | 5.60 |
| Clinical-only OLS | Ordinary least-squares regression on the clinical co-variates. | 0.3127 | 0.0435 | 4.48 | 5.63 |
| **Atlas transfer (graphTRIP weights reused unchanged)** |  |  |  |  |  |
| graphTRIP → Schaefer 200 | Model trained on Schaefer 100 applied to Schaefer 200 parcellations (Fig. 3). | 0.7156 | 0.5113 | 3.04 | 4.02 |
| graphTRIP → AAL | Model trained on Schaefer 100 applied to AAL par-cellations (Supp. Fig. 9). | 0.6049 | 0.2863 | 3.46 | 4.86 |
| **Secondary outcome** |  |  |  |  |  |
| graphTRIP (BDI outcome) | Full model retrained to predict post-treatment BDI; MAE and RMSE are in BDI points and are not com-parable to the QIDS rows (Supp. Fig. 8). | 0.5254 | 0.2390 | 6.67 | 8.72 |

**TABLE 3.** Model and training configuration for graphTRIP. The same parameters were used for training graphTRIP to predict post-treatment BDI instead of QIDS.

| Configuration | Setting |
|---|---|
| **Training configuration** |  |
| Learning rate | 0.001 |
| Number of epochs | 300 |
| Number of latent samples | 1 |
| Alpha | 0.5 |
| Batch size | 9 |
| Number of cross-validation folds | 7 |
| **MLP configuration** |  |
| Regularisation strength | 0.01 |
| Dropout probability | 0.25 |
| MLP architecture | Input: 36; Layers: [64 → 64 → 1] with LeakyReLU (slope = 0.01) and Dropout(0.25) between layers |
| **VGAE configuration** |  |
| Latent dimension | 32 |
| VGAE regularisation strength | 0.01 |
| Encoder | Graphormer-based node encoder with input projection (6 → 16), edge and shortest-path distance encodings used as additive attention biases, followed by 3 Graphormer layers (hidden dim = 16, FFN dim = 32, dropout = 0.25), and an output projection (16 → 32); variational encoding via Linear(32 → 64) to parameterise (*μ*, log *σ*<sup>2</sup>) |
| Pooling layer | GraphTransformer pooling that first applies multi-head self-attention and a feedforward network to contextualise node embeddings (32 → 32 → 32), followed by LayerNorm, Dropout(0.25), and mean pooling |
| Edge decoder | Linear(64 → 32) → LeakyReLU → Dropout(0.25) → Linear(32 → 32) → LeakyReLU → Dropout(0.25) → Linear(32 → 1); Activation: Tanh |
| Edge index decoder | Same as edge decoder; Activation: Sigmoid |
| Node decoder | Linear(32 → 32) → LeakyReLU → Dropout(0.25) → Linear(32 → 32) → LeakyReLU → Dropout(0.25) → Linear(32 → 3) |

**TABLE 4.** Model and training configuration for Medusa-graphTRIP. The causal model shares the same VGAE architecture as the core graphTRIP model, but replaces the pooling layer and prediction head to enable treatment-specific outcome estimation.

| **Configuration** | **Setting** |
|---|---|
| **Training configuration** |  |
| Learning rate | 0.001 |
| Number of epochs | 300 |
| Number of latent samples | 3 |
| Alpha | 0.5 |
| Batch size | 9 |
| Number of cross-validation folds | 7 |
| Beta (MMD loss parameter) | 1.0 |
| **CFR prediction head configuration** |  |
| Regularisation strength | 0.01 |
| Dropout probability | 0.20 |
| CFRHead architecture | Two treatment-specific RegressionMLP heads (escitalopram, psilocybin), each with Input: 67; Layers: [32 → 1] using LeakyReLU (slope = 0.01) and Dropout(0.20) |
| Treatment selection | Predictions selected via treatment indicator *t* |
| **VGAE configuration** |  |
| Latent dimension | 32 |
| VGAE regularisation strength | 0.01 |
| Encoder | Graphormer-based node encoder with input projection (6 → 16), edge and shortest-path distance encodings used as additive attention biases, followed by 3 Graphormer layers (hidden dim = 16, FFN dim = 32, dropout = 0.25), and an output projection (16 → 32); variational encoding via Linear(32 → 64) to parameterise (*μ*, log *σ*<sup>2</sup>) |
| Pooling layer | Mean–Std pooling: concatenation of per-graph mean and standard deviation of latent node embeddings (output dimension = 2 × 32) |
| Edge decoder | Linear(64 → 32) → LeakyReLU → Dropout(0.25) → Linear(32 → 32) → LeakyReLU → Dropout(0.25) → Linear(32 → 1); Activation: Tanh |
| Edge index decoder | Same as edge decoder; Activation: Sigmoid |
| Node decoder | Linear(32 → 32) → LeakyReLU → Dropout(0.25) → Linear(32 → 32) → LeakyReLU → Dropout(0.25) → Linear(32 → 3) |

### Regional attribution analysis

To quantify the contribution of individual brain regions to model predictions, we performed a regional attribution analysis based on gradients in latent space. For each patient, we computed the gradient of the model prediction with respect to the latent node representations of the brain graph. This analysis was applied both to predicted post-treatment QIDS scores (using `graphTRIP`) and to ITE estimates (using `Medusa-graphTRIP`). The resulting region-specific gradient vectors quantify the local sensitivity of the prediction to perturbations in each region’s latent representation.

Regional attribution scores were computed by first normalising the full gradient vector (obtained by concatenating all node-wise gradients) to unit Euclidean norm, and then computing, for each region, the sum of squares of its partial gradient components. This yields non-negative regional attribution scores that sum to one (or to 100 after rescaling) and can be interpreted as the percentage contribution of each region to the total gradient energy. Under small Gaussian perturbations of the latent variables, this quantity is proportional to the expected contribution of each region to the variance of the predicted outcome, consistent with established gradient-based attribution methods [37].

To obtain robust patient-level estimates, we accounted for both latent sampling variability and model stochasticity. For each patient, we sampled 100 latent vectors from a Gaussian distribution centred at the patient-specific latent mean, computed regional attribution vectors for each sample, and averaged these to obtain a stable attribution profile. We then aggregated attributions across all CV-fold models and random seeds using a performance-weighted average, with weights proportional to test-fold prediction accuracy (true–vs-predicted Spearman correlation).

### Permutation importance analysis

To quantify the contribution of each input feature to the treatment outcome predictions, we conducted permutation importance analysis. Individual features, including pre-treatment depression scores, prior SSRI use, treatment condition, and the latent brain-graph representation, were shuffled independently across patients in the test set, using 50 random permutations per feature. All other features were held constant. Feature importance was defined as the difference between the model’s mean absolute error (MAE) on the permuted input and the MAE on the original data. Larger increases in MAE indicate greater predictive importance. For the latent brain representation, the entire vector ***z*** was permuted as a unit, because permuting individual latent coordinates would yield out-of-distribution inputs for the MLP. An important limitation of this approach is that the ***z*** is 32-dimensional whereas the remaining predictors are scalars, making this a group-versus-scalar comparison.

## DATA AVAILABILITY

All requests for raw or processed data and study materials will be reviewed by R.L.C.-H., the chief investigator of both clinical trials from which the datasets were obtained. Patient-level clinical data are subject to patient confidentiality and data sharing restrictions. A minimal dataset sufficient to reproduce core figures and results is available in the supplementary materials. The normative receptor density maps from in vivo PET used in this study are publicly available: serotonin receptor and transporter maps can be accessed at https://nru.dk/index.php/allcategories/category/90-nru-serotonin-atlas-and-clustering; all other receptor and transporter maps are available at https://github.com/netneurolab/hansen_receptors.

## CODE AVAILABILITY

The code used to implement and run all analyses in this study is publicly available at: https://github.com/Imperial-MIND-lab/graphTRIP

## AUTHOR CONTRIBUTIONS

Conceptualization, H.M.T. and P.A.M.M.; Data curation, L.R. and D.N. and R.L.C.-H.; Formal analysis, H.M.T.; Investigation, H.M.T.; Methodology, H.M.T. and T.L.; Project administration, H.M.T.; Software, H.M.T. and T.L.; Supervision, P.A.M.M. and A.I.L. and R.L.C.-H.; Visualization, H.M.T.; Writing – original draft, H.M.T.; Writing – review & editing, H.M.T. and P.A.M.M. and A.I.L. and T.L.

## COMPETING INTERESTS

R.L.C.-H. is a scientific advisor to TRYP Therapeutics, MindState Design Labs, and Red Light Holland. D.N. has received lecture fees from Takeda, Lundbeck, Otsuka, and Janssen, and consulting fees from Algernon, Beckley Psytech, Leith Pharma, and Amitis Partners. He is a director of GABA Labs and Chief Research Officer at Solvonis (formerly Awaknlifesciences), and holds shares/options in PsychedWellness and Neurotherapeutics. D.N. is also Head of the Imperial College Centre for Psychedelic Research, which has received in-kind support from COMPASS Pathways and USONA (psilocy-bin), and Beckley Psytech (5-MeO-DMT). All other authors declare no competing interests.

## ACKNOWLEDGMENTS

H.M.T. is supported by the Doctoral Teaching Scholarship of the Department of Computing, Imperial College London. A.I.L. acknowledges support from St John’s College, Cambridge; and a Wellcome Early Career Award (grant number 226924/Z/23/Z). L.R. is supported by the Huxley Foundation. We are grateful to Lewis J. Ng, whose prior work on predictive modelling for this dataset provided valuable context and inspiration for this project. For the purpose of open access, the authors have applied a Creative Commons Attribution (CC-BY) license to any Author Accepted Manuscript version arising from this submission.

## Algorithm 1 graphTRIP Training Pipeline.

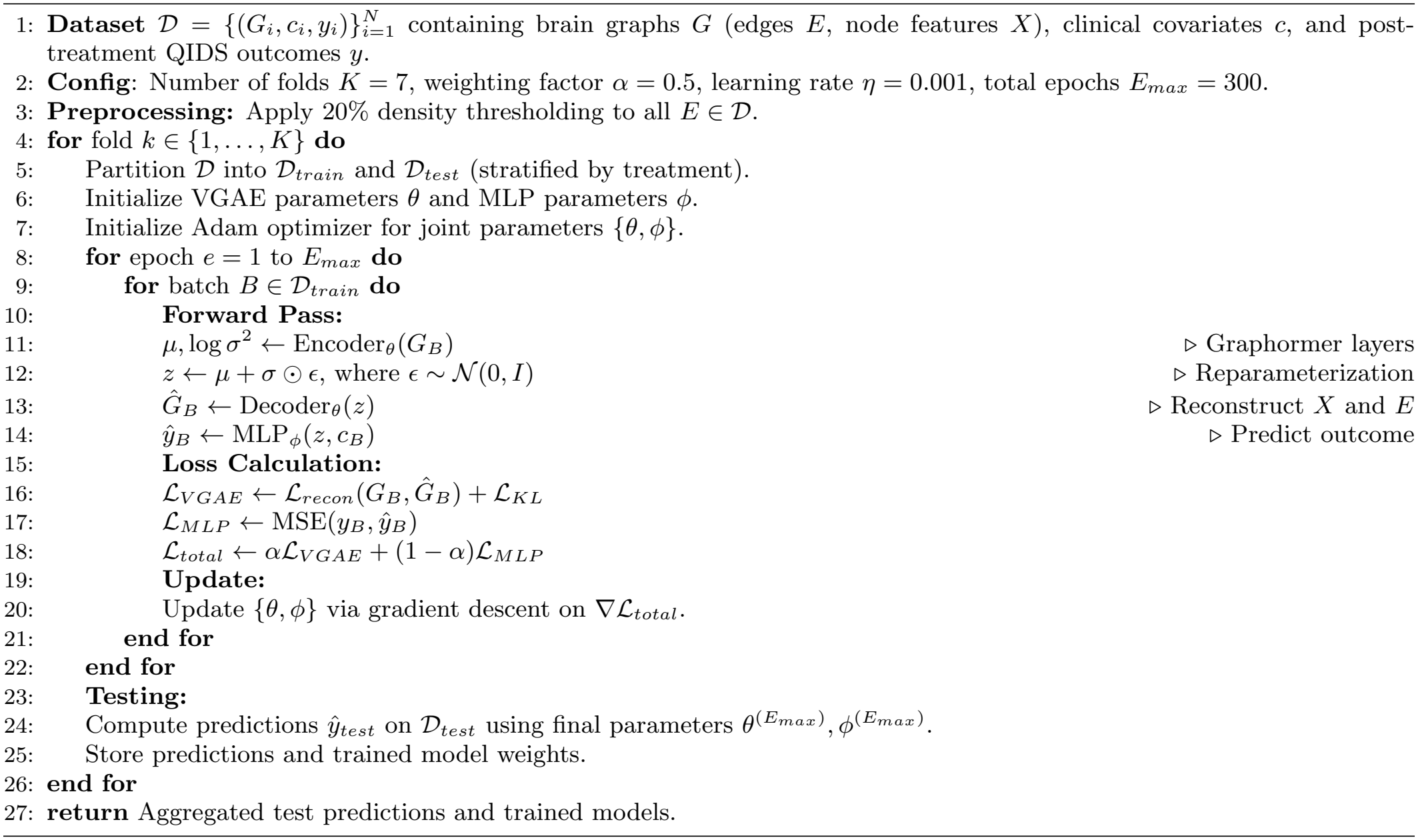

## Notes

### Summary of Updates

- reports statistical significance of model test performances using a permutation null (obtained by retraining models on shuffled labels). - revises Fig. 2 layout to structure the results into: demonstration of graphTRIP's performance and superiority over architectural ablations (a-d); and relevance of neuroimaging data inputs over clinical data inputs for the predictions (e, f). - replaces the fine-tuning approach for evaluating the model on the independent dataset (Fig. 4) by zero-shot performance without retraining. - adds a supplementary table (Tab II) listing additional performance metrics for all models in the manuscript (r, r2, mae, rmse). - adds an analysis of the variance/uncertainty of single-subject predictions (Fig. 7). - adds systematic feature ablations for graphTRIP and Medusa-graphTRIP to assess the importance of FC versus REACT features for predicting response and treatment effect (Fig. 12). - adds permutation-model based biomarker tests (Fig. 14). - adds a comparison of graphTRIP to a non-graph baseline to illustrate the effect of graphTRIP's GNN architecture in mitigating overfitting.

